# Hydrogen sulfide dynamically upregulates copper uptake and localization

**DOI:** 10.64898/2026.07.30.741779

**Authors:** Jutta Diessl, Joseph Roman, Roshan Kumar, David A. Hanna, Aaron Sue, Andrew Crawford, Romika Shokohi, Anya Parikh, Ajith Pattammattel, Andrew Kiss, Kewei Zhao, Ajay Larkin, Yibo Fu, Alex Guo, Timothy Durham, Maciek R. Antoniewicz, Si Chen, Vishal Gohil, Vamsi Mootha, Yatrik Shah, Amit R. Reddi, Kaushik Ragunathan, Ritimukta Sarangi, Thomas V. O’Halloran, Martina Ralle, Ruma Banerjee

**Affiliations:** Department of Biological Chemistry, University of Michigan Medical Center, Ann Arbor, MI; Department of Microbiology & Molecular Genetics, Michigan State University, East Lansing, MI; National Synchrotron Light Source II, Brookhaven National Laboratory, Upton, NY; SLAC National Accelerator Laboratory, Menlo Park, CA; Department of Biology, Brandeis University, Waltham, MA; School of Chemistry & Biochemistry, Georgia Institute of Technology, Atlanta, GA; Broad Institute, Cambridge, MA 02142; Department of Chemical Engineering, University of Michigan, Ann Arbor, MI; X-ray Science Division, Advanced Photon Source, Argonne National Laboratory, Lemont, IL; Department of Biochemistry and Biophysics, Texas A&M University, College Station, TX 77843; Howard Hughes Medical Institute and Department of Molecular Biology, Massachusetts General Hospital, Boston MA 02114; Department of Molecular and Integrative Physiology, University of Michigan Medical Center, Ann Arbor, MI; Department of Molecular and Medical Genetics, Oregon Health & Science University, Portland, OR

**Keywords:** Copper, hydrogen sulfide, X-ray fluorescence microscopy, X-ray absorption spectroscopy

## Abstract

The reactivity of copper, an essential micronutrient that undergoes facile cycling between Cu^1+^ and Cu^2+^ redox states, is carefully controlled within the confines of protein binding sites, and by sequestration in storage vesicles, or harnessed to kill pathogens by active pumping of Cu^1+^ into phagosomes. We have discovered that hydrogen sulfide, a signaling metabolite generated in copious quantities at the host-microbiome interface, upregulates Cu accumulation in diffusely dispersed puncta across the cell, as visualized by X-ray fluorescence microscopy. The Cu is predominantly in the Cu^2+^ state with oxygen/nitrogen ligands. Cu import occurs via the non-canonical ZNT1 transporter, while export, following sulfide withdrawal, is ATP7A-dependent. Cu accumulates at the apices of colon crypts in a mouse model of elevated sulfide exposure due to SQOR deficiency in the intestinal epithelium, establishing *in vivo* relevance. Our study reveals that sulfide is a dynamic regulator of the Cu pool, stimulating Cu^2+^ influx into highly concentrated puncta.

## INTRODUCTION

Copper is an essential micronutrient for aerobic organisms and its concentration is tightly regulated by homeostatic import/export mechanisms.^1,2^ Facile redox cycling between the cuprous (Cu^1+^) and cupric (Cu^2+^) states underlies its utility as a cofactor in processes ranging from respiration and iron homeostasis, to cell proliferation and growth. The relatively high redox potentials for free and most protein-bound Cu^2+^/Cu^1+^ couples necessitate stringent control over Cu pools to minimize spurious side reactions, including reactive oxygen species (ROS) formation via Fenton and Haber-Weiss reactions.^3^ The “static” Cu pool is tightly bound to metalloproteins with cytochrome c oxidase (or complex IV) representing a sizeable sink. In contrast, the “dynamic” pool may be transiently bound to low molecular weight ligands and allosteric sites on proteins.^4^ Important signaling roles are beginning to emerge for the dynamic Cu pool, ranging from lipolysis to sleep-wake cycle regulation.^5–8^

In yeast, the vacuole, which is similar to the mammalian lysosome, is an important compartment for Cu storage and mobilization.^9^ In brain, Cu storage vesicles that are up to 1 µm in size are found in astrocytes that are localized in the subventricular zone.^10^ The vesicles contain Cu^1+^ coordinated by sulfur ligands, resembling the multi-metallic Cu-S clusters found in metallothioneins. Sequestration in astrocytic vesicles is postulated to buffer Cu levels in the interstitial fluid^10^ and protect dendritic spines in neurons from excess Cu-dependent destabilization of F-actin.^11^ Chronic inflammation promotes Cu uptake that is driven by IL17 and NFκB activation and is associated with colon tumorigenesis.^12^ Expansion of the labile Cu pool in host epithelial cells in response to bacterial pathogens, protects against infection by activating immune signaling.^13^ As the most reactive metal in the Irving Williams series, Cu toxicity is exploited as an antimicrobial strategy within phagolysosomes.^14^ The general view that Cu^1+^ predominates in the reducing intracellular milieu, together with a paucity of tools for studying Cu^2+^, have led to a more limited understanding of how the Cu^2+^ pool is regulated.^15^ Oxidative stress due to an increase in the glutathione disulfide to glutathione ratio, or driven by oncogenic mutations, is correlated with an increase in labile Cu^2+^ and a concomitant decrease in Cu^1+^.^15^ On the other hand, deletion of NRF2, which induces antioxidant response element gene expression, increases both Cu^1+^ and Cu^2+^.^15^

The cellular redox poise, which is influenced by overnutrition, inflammation, hereditary mutations, or gas signaling agents is an important regulator of the metallome as well as the thiol proteome, a quantitatively significant redox buffer.^16,17^ Hydrogen sulfide (H_2_S) is a metabolite that, depending on its concentration, can stimulate or inhibit electron transport chain flux with pleitropic cellular and physiological consequences.^18–20^ H_2_S is derived from amino acid catabolism in host cells^21^ and its levels are sensitive to dietary intake and, importantly, microbial sulfur metabolism,^22^ which is not only responsible for high total sulfide concentrations (0.2-2.4 mM) in colon lumen^23,24, 25^ but also accounts for the majority of sulfide in circulation.^22,26^

Sulfide concentrations are maintained at very low levels in cells by its high rate of oxidation catalyzed by SQOR.^27^ When the cellular capacity for H_2_S oxidation is exceeded, profound changes in mitochondrial form and function ensue, including loss of cristae and decommissioning of O_2_-dependent energy production.^28^ The Cu-containing MT-CO1 and MT-CO2 subunits of complex IV are destabilized and energy production switches to aerobic glycolysis.^22,29,30^ Collectively, chronic sulfide increases intracellular O_2_,^31^ overreduces the NAD and CoQ pools,^32,33^ increases ROS^31^ and stimulates persulfidation,^34,35^ a protein postranslational mark that is widely believed to be important for H_2_S signaling.^36^ Persulfidation of Zn-finger proteins is associated with metal loss and disulfide formation between cysteine ligands.^37,38^

In this study, we followed up on an RNA-Seq lead that a marked decrease in metallothionein isoforms is an early response to H_2_S, and unexpectedly, discovered that sulfide induces reversible Cu accumulation across several human cell lines. X-ray fluorescence microscopy (XFM) showed that the Cu is in diffusely distributed puncta while X-ray absorption spectroscopy (XAS) revealed the predominance of Cu^2+^ with oxygen/nitrogen (O/N) ligands as expected for an intermediate Lewis acid. Consistent with Cu^2+^ accumulation, ZNT1, rather than the Cu^1+^ transporter CTR1, is involved in the sulfide response. Sulfide withdrawal leads to puncta dissolution and ATP7A- dependent Cu export. Loss of sulfide oxidation capacity in a murine model of SQOR deficiency in the intestinal epithelium, is associated with Cu elevation at crypt apices. Our study identifies sulfide as a heretofore unknown regulator of the dynamic Cu pool and establishes its physiological relevance.

## RESULTS

### Sulfide triggers metallothionein loss and Cu accumulation

An RNA-Seq analysis of colon adenocarcinoma HT-29 cells exposed to sulfide (200 µM, 2 h), revealed that the metallothionein (MT) isoforms, MT1G, MT1E, MT1F, MT1X and MT2A were significantly downregulated (Figure 1A, Supplementary Table 1). RT-qPCR analysis confirmed the decrease in MT1G, MT1X and MT2A (Supplementary Figure 1A). Since MTs bind metals, particularly Zn and Cu, we reasoned that MT downregulation would be associated with metal deficiency. However, neither pre-treatment with ZnCl_2_ nor low doses of Cu elesclomol (ES-Cu, 1- 10 nM), which preferentially replenish the mitochondrial pool,^39,40^ prevented sulfide-induced decrease in MT1G (Supplementary Figure 1B,C). In contrast, high ES-Cu (100 nM), which replenishes extra-mitochondrial stores,^39^ suppressed MT downregulation (Supplementary Figure 1C). Whole cell elemental analysis via total reflection X-ray fluorescence (TXRF) analysis showed that the Fe, Cu and Zn pools were unaffected by acute sulfide exposure (Supplementary Figure 1D). Since H_2_S disappears with a t^1^/_2_ of 3-4 min under cell culture conditions leading to experimental variability,^30^ the effect of chronic, low level exposure using a sulfide growth chamber^41^ was examined next. Unless specified otherwise, chronic exposure refers to 100 ppm H_2_S (or ∼20 µM dissolved sulfide) for 24 h.

**Figure 1.**
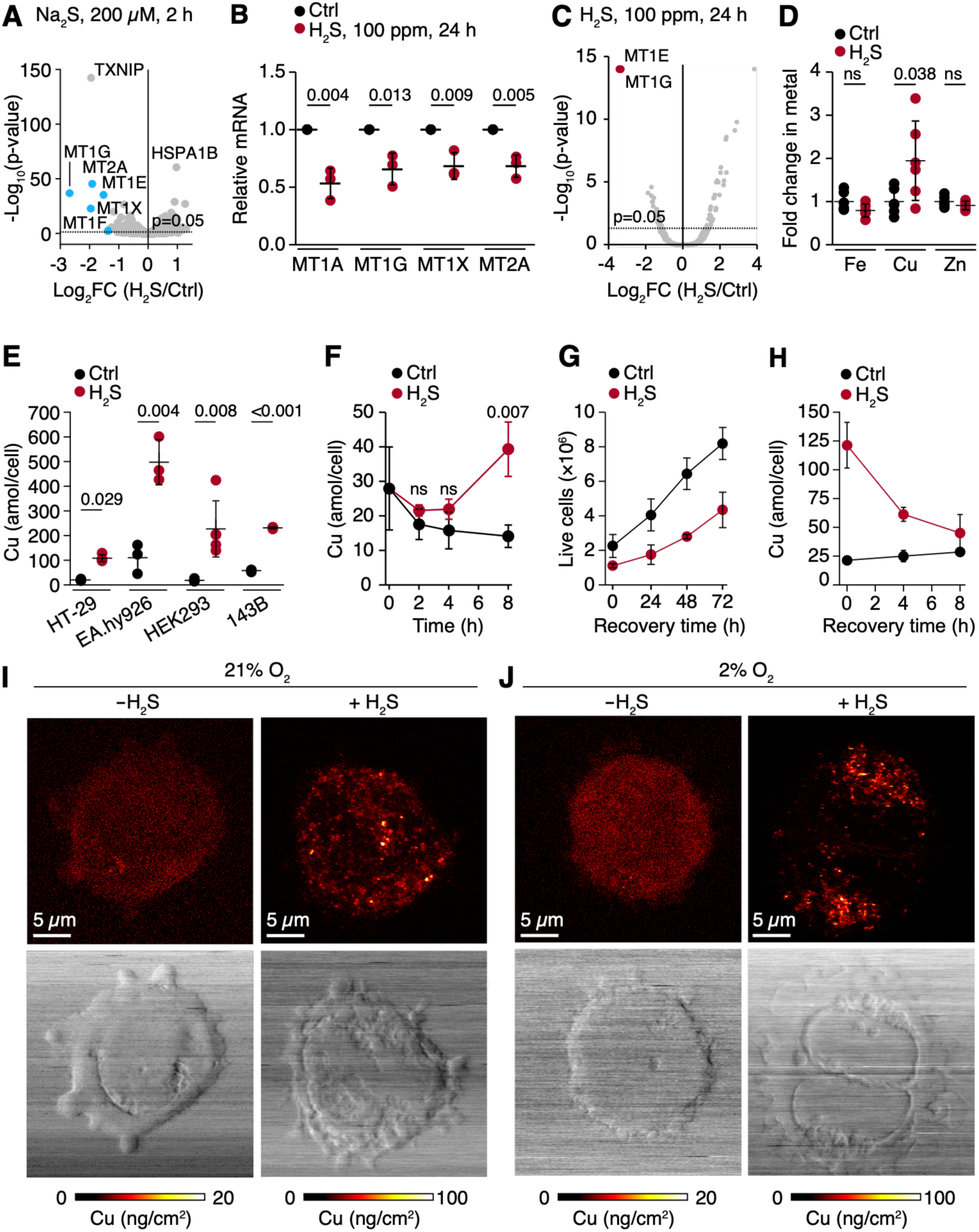
Sulfide exposure increases intracellular Cu puncta. **A.** RNA-Seq analysis reveals significant downregulation of MT1G, MT1X and MT2A in HT-29 cells in response to H_2_S (200 µM, 2 h). **B-D**. Chronic H_2_S (100 ppm, 24 h) decreases MT1A/G/X and MT2A mRNA (B), significantly lowers MT1G/E protein as seen by TMT proteomics (C), and increases Cu, but not Fe or Zn as assessed by TXRF (D). **E,F**. Sulfide increases Cu across human cell lines grown at 21% O_2_ as measured by ICP-MS. (**F-H**) Cu accumulation is detectable at 8 h following chronic H_2_S exposure in HT-29 cells (F), and cells resume proliferation following H_2_S withdrawal (F) while Cu levels normalize within 8 h (H). The data represent mean ± SD of at least 3 independent experiments in B and E-H. In D, data show mean ± SD from 2 independent experiments; each point is a technical replicate. **I,J.** Representative XFM images of HT-29 cells exposed to H_2_S in the presence of 21% (I) % or 2% O_2_ (J), showing the appearance of Cu puncta (*upper*) and the corresponding phase contrast images (*lower*).

MT1A, MT1G, MT1X and MT2A mRNA were decreased by ∼30-50% in HT-29 cells after chronic H_2_S exposure (Figure 1B). Quantitative TMT proteomics revealed an ∼11-fold decrease in MT1G/E, which could not be distinguished based on the detected peptides (Figure 1C, Supplementary Table 2). TXRF analysis revealed an increase in Cu, which was unexpected given the decrease in metal-storing MTs; Zn and Fe levels were however, unchanged (Figure 1D). We note that while Fe and Zn estimates were within a narrow range across biological replicates, a significant variation in the Cu pool was consistently seen in this and in other measurements as well. Inductively coupled plasma mass spectrometry (ICP-MS) confirmed sulfide-induced Cu accumulation across cell lines, including HT-29 (5-fold), EA.hy926 (6-fold), HEK293 (13-fold) and 143B (4-fold) (Figure 1E). In HT-29 cells, Cu accumulation could be detected reliably after 8 h (Figure 1F). As reported previously, chronic H_2_S exposure induces S-phase arrest,^28^ but following sulfide withdrawal, cells resumed proliferation after a lag, and Cu levels normalized (Figure 1G,H, and Supplementary Figure 2A,B).

### Sulfide induces Cu puncta formation

We used XFM to visualize Cu in HT-29 and HEK293 cells chronically exposed to H_2_S. Cu accumulation was readily visualized as puncta that were 300-400 nm in diameter and similar in both cell lines (Figure 1I and Supplementary Figure 2C). The diffuse distribution of Cu puncta suggested cytoplasmic localization. As colonocytes are typically exposed to low luminal O_2_, Cu puncta were also visualized in HT-29 grown in 2% O_2_ (with 100 ppm H_2_S, 24 h) and found to be comparable to the 21% O_2_ data (Figure 1I,J). Chronic H_2_S also increased Cu levels ∼5-fold in hypoxically grown HT-29 cells (from 19.9 ± 2.4 to 91.4 ± 26.2 amol/cell), indicating that H_2_S rather than low O_2_, induced Cu accumulation.

### Cu accumulation is independent of CTR1

CTR1 is a high-affinity Cu uptake protein that preferentially imports Cu^1+^ (Figure 2A).^42^ Inefficient knockdown (KD) of CTR1 in HT-29 cells was achieved with two shRNA sequences resulting in 20% and 50% decrease in protein expression, respectively (Supplementary Figure 3A). Steady-state Cu levels decreased in parallel (∼20 and 57%) in the two CTR1 KD lines (Supplementary Figure 3B) but H_2_S-induced Cu accumulation was unaffected (Figure 2B). CTR1 involvement was further investigated in H92c cells in which efficient CRISPR KO of CTR1 has been reported previously.^43^ In comparison to control H92c cells, where a 10-fold increase in Cu was seen in response to chronic sulfide, Cu levels increased 66-fold in CTR1 KOs (Figure 2C), ruling out the involvement of CTR1 in sulfide-induced Cu import. As expected, extracellular Cu chelation by bathocuproine disulfonate (BCS) abrogated Cu accumulation (Figure 2D).

**Figure 2.**
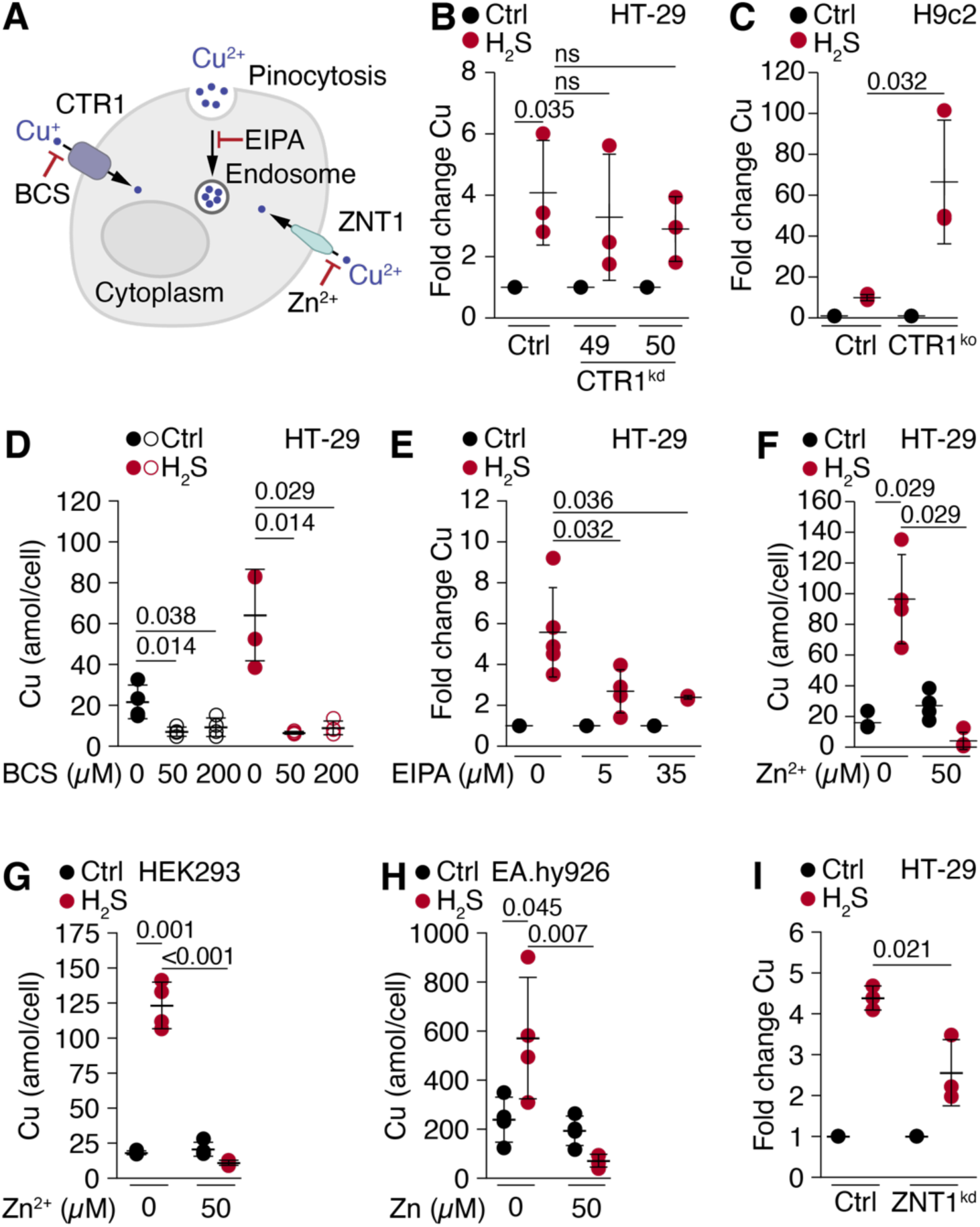
Zn prevents Cu accumulation in response to chronic H_2_S. **A.** Scheme showing Cu import pathways. **B**. CTR1 shRNA KD has a statistically insignificant effect on Cu accumulation in HT-29 cells. **C**. CRISPR KO of CTR1 in H9c2 cells leads to Cu hyperaccumulation in response to chronic H_2_S compared to control cells. **D,E.** Chelation by BCS prevents Cu accumulation (E) while inhibition of macropinocytosis by EIPA attenuates Cu accumulation (E) in HT-29 cells. **F-H**, Zn^2+^ (50 µM) prevents sulfide-induced Cu accumulation in HT-29 (F), HEK293 (G) and EA.hy926 (H) cells. **I.** ZNT1 KD attenuates sulfide-induced Cu accumulation. The data represent mean ± SD of at least 3 independent experiments in B-I.

### Cu accumulation is prevented by Zn

Next, we tested macropinocytosis^44^ and the zinc exporter 1/Cu importer ZNT1^45^ as alternative routes for Cu entry (Figure 2A). The macropinocytosis inhibitor 5-(*N*-ethyl-*N*-isopropyl)amiloride (EIPA), which targets the Na^+^/H^+^ exchanger, decreased Cu accumulation ∼50% across a 7-fold concentration range beyond which it was toxic to HT-29 cells (Figure 2E). Zn supplementation inhibits ZNT1-dependent Cu uptake,^45^ and completely inhibited Cu accumulation in response to H_2_S in all three cell lines that were tested (Figures 2F-H). ZNT1 KD efficiency with two guide sequences was evaluated by ICP-MS analysis due to the poor quality of the commercially available anti-ZNT1 antibodies. The two ZNT1 KD lines exhibited ∼1.3-fold higher basal Zn compared to control cells (Supplementary Figure 3C), signaling low KD efficiency likely due to the essentiality of ZNT1.^46^ The sg4 line was selected for further analysis and showed a ∼2-fold decrease in Cu accumulation compared to the CRISPR OR7G3 non-targeting control (Figure 2I).

### Cu puncta formation is independent of ATP7A

Since intracellular Cu returns to basal levels following H_2_S withdrawal (Figure 1H), we examined involvement of the canonical Cu transporter ATP7A. ATP7A pumps excess Cu into Golgi-derived vesicles destined for exocytosis or exports Cu directly at the plasma membrane^47^ (Figure 3A). Interestingly, ATP7A levels increased ∼1.5-fold in response to chronic sulfide (Figures 3B,C). To assess whether the puncta observed by XFM represent Cu-rich vesicles loaded by ATP7A, we tracked its co-localization with giantin, a Golgi marker protein. Sulfide did not however, affect the degree of co-localization of the two proteins (Figures 3D,E) as would be expected if ATP7A relocated from the Golgi apparatus to vesicles or the plasma membrane. Further, a difference in the number of endosomes was not observed by transmission electron microscopy analysis of HT-29 cells exposed to chronic sulfide (Supplementary Figure 4A-C). The area and perimeter of the endosomes were very similar although a wider distribution was observed in untreated cells (Supplementary Figure 4D,E).

**Figure 3.**
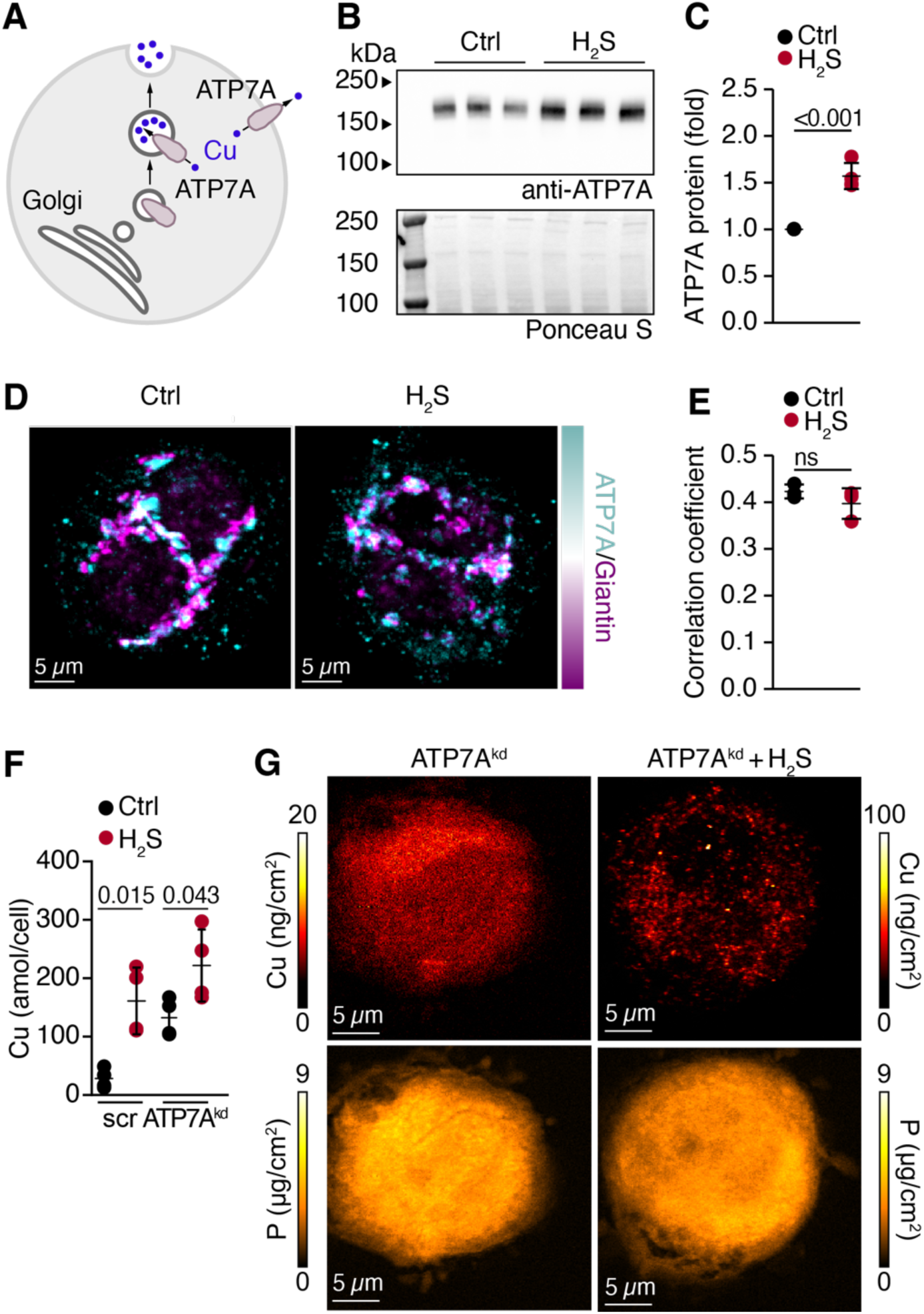
ATP7A is not required for Cu puncta formation. **A.** Scheme showing role of ATP7A in Cu export via exocytosis or pumping at the plasma membrane. **B-E**. Sulfide increases ATP7A protein in HT-29 cells (B) as quantified by western blot analysis (C, n=3) but does not change its co-localization with the Golgi protein giantin (D) as visualized by immunohistochemistry and quantified by the Pearson correlation coefficient, representing the spatial overlap of ATP7A and giantin, where a value of 1 indicates complete co-localization (E, n=3). **F,G**. ATP7A KD in HT-29 cells attenuates sulfide-induced Cu accumulation (F, n=4) but does not affect Cu puncta formation as visualized by XFM (G). Representative images are shown in D and G.

Three shRNA sequences were used to generate ATP7A KD in HT-29 cells, and one (#73), exhibiting the highest efficiency (96%), was characterized further (Supplementary Figure 3D). ATP7A KD of the primary Cu exporter increased basal Cu levels (∼5-fold) as expected, but attenuated Cu accumulation in response to H_2_S from ∼6-fold in scrambled controls to ∼2-fold in ATP7A KD cells (Figure 3F). Interestingly, loss of ATP7A did not inhibit puncta formation (Figure 3G).

In contrast to Cu puncta formation, ATP7A KD impeded recovery of basal Cu levels even 24 h after sulfide withdrawal (Supplementary Figure 5A-C). While Cu remained elevated under these conditions, Cu puncta were no longer visible, indicating that ATP7A is required to export Cu during puncta dissolution.

### Cu^2+^ with N/O ligands predominates in sulfide-treated cells

Whole cell Cu K-edge XAS and extended X-ray absorption fine structure (EXAFS) spectroscopy analyses were performed to probe the Cu oxidation state and ligand environment. Cu Kα X-ray analysis of HT-29 cells grown with chronic H_2_S exposure revealed Cu accumulation signaled by ∼10,000-13,000 fluorescence counts in the selected energy window versus ∼1,000- 2,000 counts in untreated cells. Comparison of the Cu K-edge spectrum with Cu standards allowed assignment of the rising-edge features at 8982 and 8986 eV, corresponding to dipole allowed 1s→4p transitions, to Cu^1+^ and Cu^2+^, respectively, and the pre-edge feature at ∼8979 eV, corresponding to the dipole forbidden 1s→3d transition, to Cu^2+^ (Figure 4A,B and Supplementary Figure 6A,B). Spectra were measured at 11 spots in whole cell samples, with a ∼250 µm spot size. The spectra displayed heterogeneity, both in the total amount of Cu and the chemical nature of the Cu species present in the various spots. Heterogeneity in the total amount of Cu was expected, since the sample itself is not homogeneous and the spots with fewer cells lead to lower overall Cu signal. For those spots, a weak background Cu signal was found to mix into and confound the experimental data and they were excluded from the speciation analysis. The variability between spots is shown in Supplementary Figure 6C. Linear combination fitting estimated that Cu^2+^ represents ∼75-85% and Cu^1+^ ∼15-25% of the expanded Cu pool. Since some photoreduction was observed in subsequent scans (Supplementary Figure 6D), the Cu^1+^ estimates should be considered the upper limit. The white line at ∼8998 eV is consistent with N/O- ligated Cu standards. In agreement, best fits of the EXAFS data were obtained with light atoms in the first coordination-shell; specifically three N/O ligands at 1.96 Å and one weaker N/O ligand at 2.52 Å (Figure 4C, Supplementary Table 3). This longer component may reflect a water or other weak light atom ligand. The absence of Cu–S scattering (even at a fractional contribution of 0.3 paths) strongly argues against direct sulfide or protein-based thiolate coordination. It is likely, however, that a smaller fraction of reduced Cu^1+^ is coordinated to sulfidic ligands (≤0.15 Cu–S components), which would not be discernible from the EXAFS data. Since the number and proportion of Cu species are not known, the value of the first shell EXAFS analysis is to directionally suggest the type of accumulated species rather than to specifically identify the ligand set based on the first shell bond distances.

**Figure 4.**
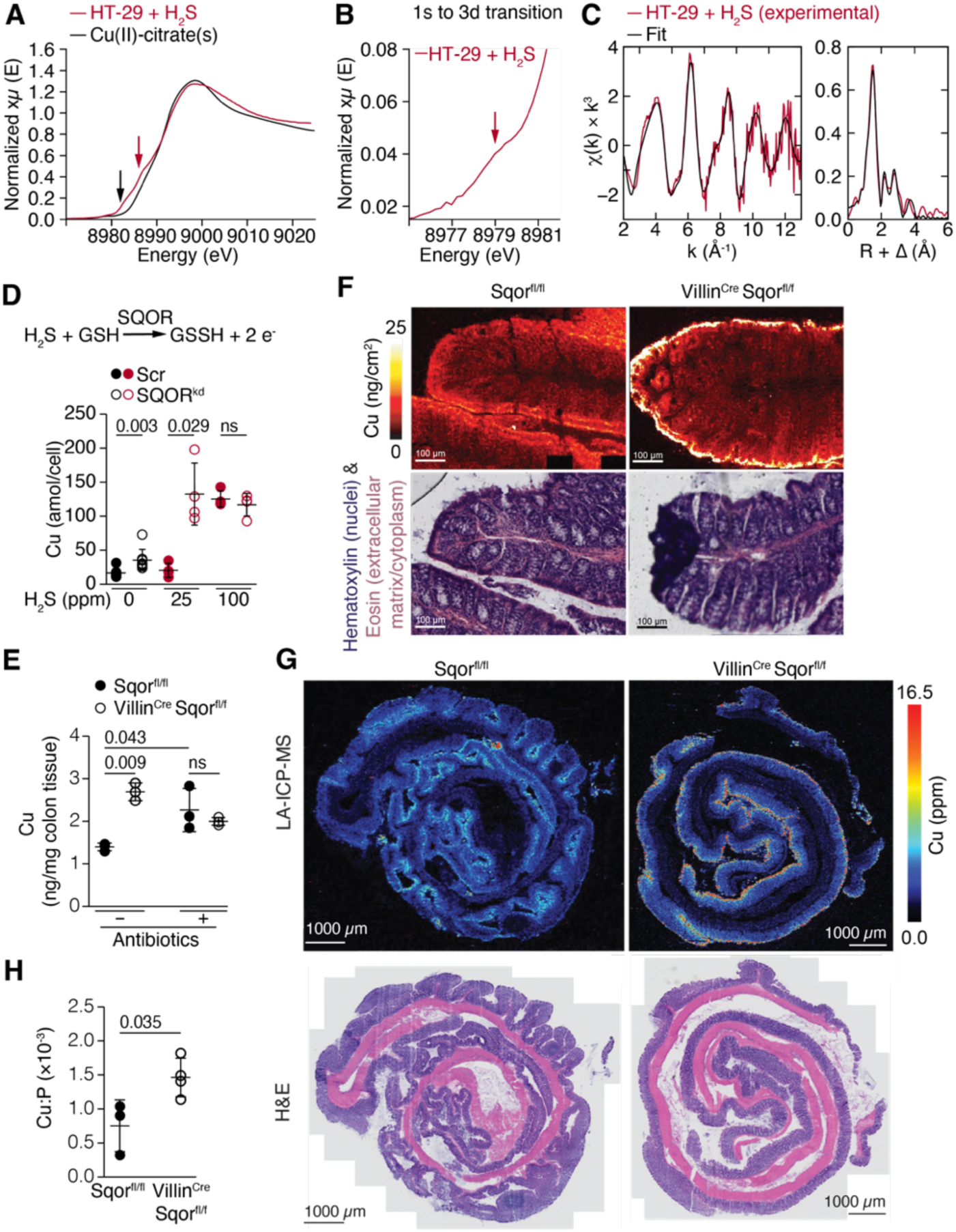
Loss of H_2_S oxidation capacity leads to Cu accumulation in the form of Cu^2+^ with N/O ligands. **A, B**. XAS reveals that Cu the predominance of Cu^2+^ as indicated by the rising-edge feature at 8986 eV (A, red arrow) and a pre-edge feature at 8979 eV (B, red arrow) while Cu^1+^ represents a minor fraction as indicated by a rising-edge feature at 8982 eV (black arrow). **C.** EXAFS fitting reveals that the first-shell ligands are light atoms (N/O). Left: Experimental k^3^- weighted EXAFS oscillations χ(k) (red line) and the corresponding best-fit model (black line). Right: Fourier-transformed magnitude of the k^3^-weighted data (solid black line) and the best-fit model (dashed red line) in R-space. **D**. SQOR KD cells show higher basal Cu and increased sensitivity to H_2_S-induced Cu accumulation at low H_2_S (25 ppm, 24 h) compared to scrambled control (Scr) HT-29 cells (n=4). **E-H**. Villin^Cre^ SQOR^fl/fl^ mice show increased Cu in colon tissue quantified via ICP-MS (E, n=3), which is localized at the apices of colon crypts as seen by XFM (F) and LA-ICP-MS imaging (G). Quantitative analysis of Cu in G is shown in H (n=3). Hematoxylin and Eosin (H&E) staining of an adjacent colon section used for XFM (F) or adjacent cross section of the Swiss roll used for LA-ICP-MS imaging (H) is shown below the respective representative images.

The presence of outer-shell scattering features in the EXAFS Fourier transform strongly indicates the presence of a dominant Cu^2+^ component with chelating ligands such as citrate. If, on the other hand, the accumulation of Cu was highly heterogeneous, the EXAFS data would be expected to decay more rapidly, without producing the higher R features in the Fourier transform. This prompted us to consider whether a low molecular weight metabolite might serve as a Cu^2+^ ligand. Examination of published metabolomics analysis of HT-29 exposed to H_2_S revealed carbamoyl aspartate (CA) accumulation,^29^ which was validated by GC-MS to increase 175-fold from 6 ± 2 to 1048 ± 334 µM (Supplementary Figure 7A). CA is an intermediate in the pyrimidine biosynthetic pathway and upstream of dihydroorotate, which also accumulates due to overreduction of the CoQ pool but being cyclic, is less likely to chelate Cu (Supplementary Figure 7B,C). In comparison to citrate, which is a 6-carbon compound with three carboxylates, CA is an O/N rich 5-carbon compound with two carboxylates and a carbamate group. To assess the feasibility of CA serving as a Cu ligand, heterologous expression of *Lactobacillus brevis* NADH oxidase (*Lb*NOX) was used to alleviate the sulfide-induced reductive shift (Supplementary Figure 7C). Mitochondrial but not cytosolic expression of *Lb*NOX decreased CA 7-fold (from 1312 ± 147 to 190 ± 55 µM) in sulfide-treated cells (Supplementary Figure 7A), but did not affect Cu accumulation (Supplementary Figure 7D). Further, while basal Cu levels were seen within 8 h of sulfide withdrawal (Figure 1H), CA levels remained elevated at this time point (Supplementary Figure 7E). These data are not consistent with the involvement of CA as a Cu^2+^ ligand. Levels of CAD, the trifunctional enzyme, which harbors aspartate transcarbamylase activity and catalyzes the synthesis of CA, were decreased ∼60% by two of the three shRNA sequences (Supplementary Figure 7F). XFM analysis of scrambled control versus CAD KD HT-29 cells revealed no differences in Cu puncta formation in response to chronic H_2_S exposure Supplementary Figure 7G).

### Decreased H₂S clearance increases cellular and tissue Cu

Since SQOR catalyzes the two-electron oxidation of H_2_S and is critical for maintaining low intracellular levels of sulfide,^27^ its deficiency is predicted to increase Cu levels. Indeed, basal Cu levels were two-fold higher in SQOR KD HT-29 cells (35 ± 16 amol/cell) compared to scrambled controls (17 ± 6 amol/cell) (Figure 4D). Further, SQOR KD cells were more sensitive to sulfide and showed a 4-fold increase in Cu with 25 ppm H_2_S (∼4 µM dissolved sulfide) exposure while a comparable increase was observed in control cells only at 100 ppm H_2_S (Figure 4D).

The *in vivo* relevance of Cu accumulation in response to sulfide was assessed in the *Villin*^Cre^ SQOR^fl/fl^ mouse model,^22^ lacking SQOR in the intestinal epithelium.^22^ ICP-MS analysis revealed 2-fold higher Cu in colon from *Villin*^Cre^ SQOR^fl/fl^ versus control SQOR^fl/fl^ mice (Figure 4E), which was concentrated at the crypt apices as revealed by XFM analysis (Figure 4F). Apical localization and an increase in Cu content were also observed by laser ablation ICP-MS in colon from *Villin*^Cre^ SQOR^fl/fl^ mice (Figure 4G,H). Antibiotics dramatically decrease gut sulfide exposure since microbes are the dominant source of sulfide.^22^ Interestingly, while basal Cu levels were higher in control mice, presumably due to decreased competition from microbes, loss of sulfide oxidation capacity in colon of *Villin*^Cre^ SQOR^fl/fl^ mice had no further effect on Cu levels compared to controls (Figure 4E).

## DISCUSSION

Sulfide toxicity is averted by efficient SQOR-dependent oxidation at the electron transport chain that relies on Cu-dependent cytochrome *c* oxidase and O_2_ as a terminal electron acceptor. When the host capacity to shield against sulfide toxicity is exceeded, cells switch to a low respiratory mode characterized by remodeled O_2_, lipid and amino acid metabolism.^19,20^ In this study, we have uncovered Cu uptake and sequestration as another facet of this survival phenotype. We find that Cu uptake is mediated by a pathway that is distinct from the Cu^1+^- dependent antimicrobial strategies described for immune cells and inflamed tissues.^12,14,48^

H_2_S is slightly hydrophobic, which promotes both its concentration in and unfacilitated permeation across cell membranes.^49,50^ Once inside, H_2_S rapidly dissociates, releasing a proton and the sulfide anion. At pH 7.4, the proportion of HS^-^ and H_2_S are 81 and 19%, respectively,^18^ and chronic exposure is predicted to lead to sulfide accumulation and acidification (Figure 5). An H_2_S dose dependent decrease in intracellular pH has been reported previously, but attributed to inhibition of the Na^+^/H^+^ exchanger.^51^ We speculate that the observed destabilization of MTs results from loss of Cu due to protonation of thiolate ligands and increased proteolytic susceptibility of the resulting apo-proteins. Sulfide sensitivity is restricted to a subset of isoforms (MT1G, MT1X and MT2A) and a decrease in their mRNAs levels is an early response (Figure 1A). The mechanism underlying the selective loss of MT mRNA and protein isoforms is presently unknown. Paradoxically, MT loss is accompanied by a dynamic expansion predominantly of the Cu^2+^ pool, and its accumulation in puncta, which, based on the size and diffuse distribution, is likely to be vesicular (Figure 1). Cu accumulation in an acidic lysosome-related organelle, previously refered to as acidocalcisome, in response to Zn^2+^ deficiency has been seen in *Chlamydomonas reinhardtii*.^52^ The Cu is however, predominantly Cu^1+^ with a sulfur ligand, presumed to be derived from cysteine.^53^

**Figure 5.**
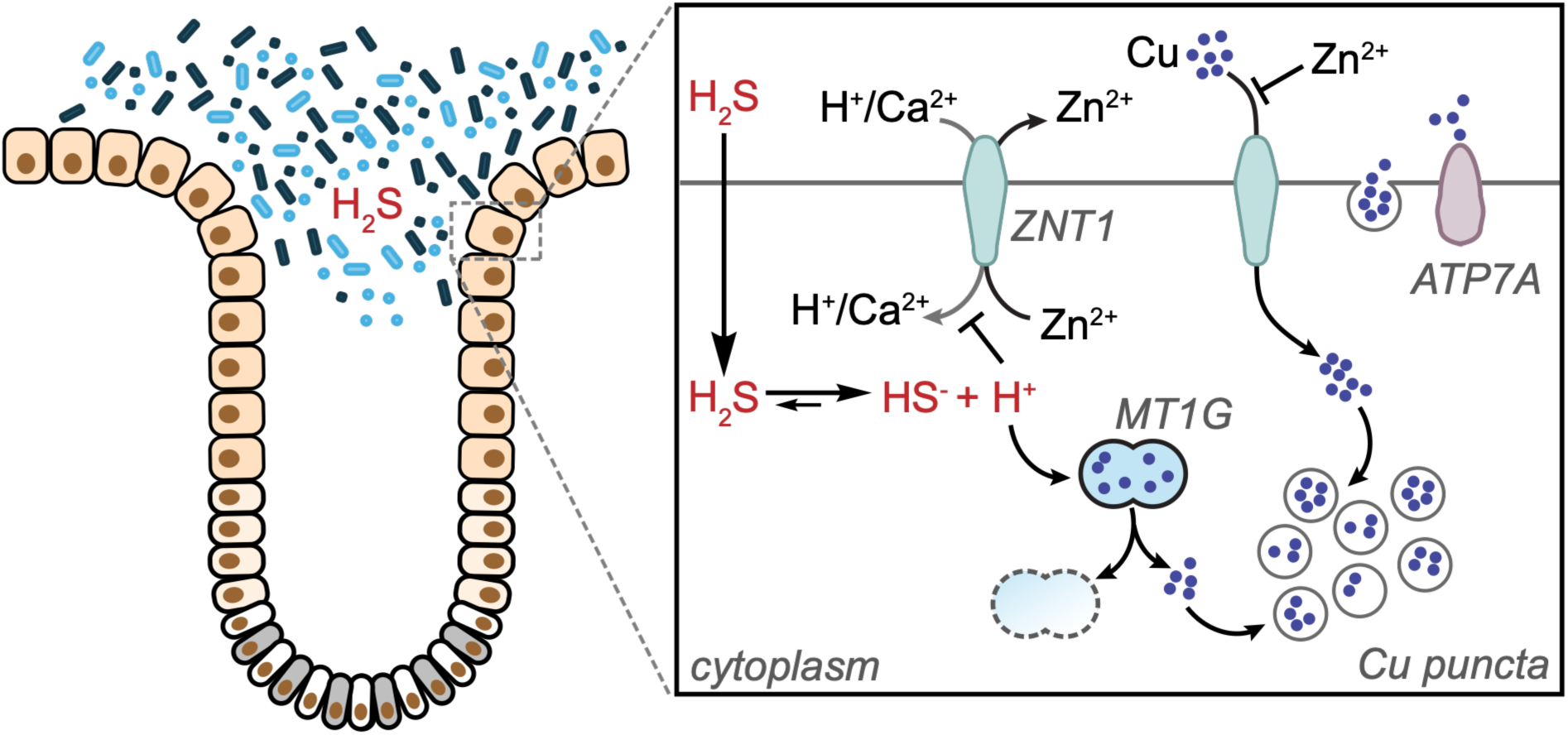
Model for dynamic regulation of the Cu pool by H_2_S. Colonocytes form a single-layered barrier between host and microbes (blue and black rods and spheres) with stem cells located at the bottom of crypts (*left*). Close-up of H_2_S-driven intracellular changes affecting the Cu pool (*right).* Unfacilitated permeation of H_2_S across the plasma membrane is followed by rapid pH-sensitive dissociation, leading to acidification and a proposed loss of Cu from select metallothionein isoforms, ZNT1-dependent Cu^2+^ import (which is inhibited by extracellular Zn^2+^), and sequestration of Cu in puncta. Following H_2_S withdrawal, puncta disappear and Cu extrusion is dependent on ATP7A.

Our data suggest a primary role for ZNT1 in sulfide-activated Cu^2+^ import. ZNT1 is the principal Zn exporter at the plasma membrane and its activity is coupled to either Ca^2+^ or H^+^ import.^46,54^ An increase in the intracellular H^+^ concentration due to H_2_S dissociation at physiological pH is predicted to inhibit Zn export, which we speculate activates Cu^2+^ import (Figure 5). In the cytoplasmic domain, protonation of the histidine and glutamate residues that serve as Zn ligands in ZNT1 would weaken binding. In the extracellular domain, an inter-subunit disulfide is needed for efficient Cu import, suggesting susceptibility to redox regulation.^45^ Our data do not however, support a model of ZNT1 persulfidation via nucleophilic addition of the sulfide anion to the disulfide bond as an activation mechanism for Cu import. While Zn supplementation increased MT mRNA as expected, sulfide still induced a decrease in MT1G levels (Supplementary Figure 1B), making it unlikely that the Zn effect on Cu was due to MT stabilization. Unlike Zn, which completely inhibited Cu accumulation, EIPA, decreased it by 50 percent (Figure 2E,F). In addition to inhibiting the Na^+^/H^+^ exchanger, EIPA has off-target effects on F-actin reorganization and endosome morphology and distribution,^55^ which could confound its effects on Cu import. Alternatively, since ZNT1 can also localize to endosomes,^56^ the inhibitory effect of EIPA might be downstream of Cu import. While ATP7A is not involved in Cu loading into puncta, it is needed for Cu extrusion following H_2_S withdrawal (Figure 3G).

Sulfide-driven expansion of the dynamic Cu pool provides new insights into its regulation but also raises a number of questions. The involvement of a subset of MT isoforms suggests a functional division of labor, namely that MT1G, MT1X and MT2A are chiefly involved in Cu buffering. What is the mechanism by which mRNAs encoding specific MT isoforms are downregulated, and is mitochondrial signaling involved in cytosolic puncta formation? Boosting the cytoplasmic but not the mitochondrial Cu pool with ES-Cu suppressed sulfide-dependent downregulation of MT1G mRNA (Supplementary Figure 1C), indicating that at least this early response is extramitochondrial. It is unknown whether the Cu ligands in puncta are low molecular weight metabolites, peptides or proteins. If Cu accumulation occurs in response to acidification resulting from rapid pH-driven equilibration of dissolved H_2_S, is alkalinization involved in puncta dissolution and Cu extrusion? Finally, what is the physiological significance of sulfide-sensitive Cu accumulation?

In the gut, the epithelium represents a single-cell layered frontier for the interaction between self and microbiota, and is teeming with a wealth of metabolites. An estimated two-thirds of systemic host sulfide levels is attributed to microbial metabolism.^22,26^ One possibility is that chronically elevated H_2_S is read as a danger signal and activates an epithelial response that includes Cu sequestration as a defensive nutritional immunity strategy. Alternatively, Cu sequestration might represent a strategy for averting precipitation as a sulfide salt; the solubility products for Cu^1+^ (2 x 10^-47^) and Cu^2+^ (6 x 10^-36^) sulfides are among the lowest for biologically relevant metals.^57^ As the “threat” subsides, i.e. sulfide is withdrawn, cells extrude Cu.

In summary, we have identified sulfide as a previously unknown regulator of dynamic Cu pools, which induces reversible sequestration of Cu^2+^ as part of a larger metabolic adaptation to a stress signal. An important implication of sulfide-induced mitochondrial remodeling and Cu sequestration is that it protects cells from cuproptosis, which depends on respiratory metabolism and active pyruvate flux into the TCA cycle.^58^

## Data availability

All data generated and analyzed in this study are included in the main text and Supplementary Information file. The RNA-Seq data (Table S1) have been uploaded in the GEO database (accession number: GSE341519) and the TMT-proteomics data (Table S2) have been deposited in the Pride repository (dataset identifier: PXD079024).

## Acknowledgements

This work was supported in part by the grants from the National Institutes of Health (R35GM130183 to RB, F32GM154404 to JD, R35GM152102 to VMG, R35GM145350 to ARR, P41GM135018, R01GM115848, and R01GM038784 to TVO). We thank Dr. Einar Ólafsson (University of Michigan) for his assistance with microscopy image analysis, Dr. Antentor Hinton (Vanderbilt University) for help with TEM image analysis, and Dr. Yang Yang (Brookhaven National Laboratory) for advice on tissue XFM analysis. This research used the following U.S. Department of Energy (DOE) Office of Science User Facilities: the Advanced Photon Source at Argonne National Laboratory (Contract DE-AC02-06CH11357); the HXN (3-ID) and SRX (5-ID) beamlines at the National Synchrotron Light Source II, Brookhaven National Laboratory (Contract DE- SC0012704); and the Stanford Synchrotron Radiation Lightsource, SLAC National Accelerator Laboratory (Contract DE-AC02-76SF00515). The SSRL Structural Molecular Biology Program is supported by the DOE Office of Biological and Environmental Research and NIH, National Institute of General Medical Sciences (P30GM133894). Content is solely the authors’ responsibility and does not necessarily represent official NIH views.

## Author contributions

JD and RB conceptualized the study and JD performed and analyzed the majority of the experiments and was assisted by: RS and AP-Western blot, cell proliferation and ICP MS analyses; RK and YS-mouse studies, DH-TEM analysis; DH, JR, AS, AC, AK, SC, AP, TVO and MR-XFM analysis, AS, AC, TVO-LA ICP MS, AL, RK, AG, TD, VM and KR-RNA-Seq data generation and analysis; KZ and RS-XAS data; YF and ARR-TXRF, VMG-provision of ES-Cu and H92c cells, early experimental design and data analysis; and MRA-GC MS analysis. JD and RB drafted the manuscript and all authors edited and approved the final version.

## Competing Financial Interests

RB is a consultant for Zyphore Therapeutics.

