## Supplementary Materials for "Hydrogen sulfide dynamically upregulates copper uptake and localization"

**Table of Content**

Supplementary Table 1. RNA-Seq data for HT-29 cells exposed to $H_2S$
Supplementary Table 2: TMT proteomics data for HT-29 cells exposed to $H_2S$
Supplementary Table 3. Cu K-edge EXAFS curve-fitting analysis
Supplementary Table 4. pLKO.1 shRNA vectors
Supplementary Table 5. Oligonucleotides used for qPCR
Supplementary Figure 1. Acute $H_2S$ exposure decreases MT but not intracellular Cu
Supplementary Figure 2. Sulfide induces Cu puncta formation in HEK293 cells.
Supplementary Figure 3. Validation of CTR1, ATP7A and ZNT1 KDs in HT-29 cells
Supplementary Figure 4. Transmission electron microscopy analysis of endosomes in HT-29 cells
Supplementary Figure 5. ATP7A is required for Cu export, but not dissolution of Cu puncta after sulfide withdrawal
Supplementary Figure 6. Effects of spot-to-spot variation and sample photoreduction on Cu K-edge XAS
Supplementary Figure 7. $H_2S$-induced Cu accumulation is independent of carbamoyl aspartate.

### METHODS

#### Materials

**Cell lines.** HT-29 (human colorectal adenocarcinoma), EA.hy926 (human endothelial), and HEK293 (human embryonic kidney) cells were obtained from the American Type Culture Collection (ATCC). 143B (human osteosarcoma) cells were a gift from Dr. Matthew Vander Heiden (MIT). The generation of Ctr1 KO in H9c2 (rat cardiomyocyte) cells[1] and SQOR KD in HT-29 cells[2] have been described previously.

**Reagents.** Bathocuproine disulfonate (BCS, B1125), DMSO (D2653), hydrogen peroxide (31642), poly-L-lysine (P4707), Ponceau S (P3504), protease inhibitor cocktail (P8340), puromycin (P8833), sodium sulfide nonahydrate (431648), uridine (U3003), $ZnSO_4$ (Z0251), pLKO.1 shRNA vectors (TRCN0000043349, TRCN0000043350 for CTR1, TRCN0000043173, TRCN0000043174, TRCN0000043177 for ATP7A, TRCN0000045908, TRCN0000045910, TRCN0000045912 for CAD), Hoechst 33342 (AMBH9A260690) and derivatization reagent MTBSTFA + TBDMCS (375934) were from Sigma. Geneticin/G418 (10131035), DMEM (11995065), fetal bovine serum (FBS, Gibco, A5256701), HEPES (22400105), Opti-MEM (31985062), PBS (10010023), penicillin–streptomycin (P/S, 15140122), RPMI 1640 (11875093), RPMI 1640 medium with 25 mM HEPES (22400105) were from Gibco. dNTP mix (18427013), lipofectamine 3000 (L3000008) and P3000 reagent (L3000001), M-MLV Reverse Transcriptase (28025-013), random hexamers (N8080127), TRIzol (15596026), Trypan Blue (15250061), HCS CellMask DeepRed (H32721), anti-rabbit Alexa Fluor™ 555 (A-31572), anti-mouse Alexa Fluor™ 488 (R37114) were from Invitrogen. ECL substrate (1705061), Bradford reagent (5000006), PVDF membranes (0.2 µm, 1620177), Trans-Blot Turbo Transfer Kit (1704272) were from Bio-Rad. psPAX2 (12260), pLentiCRISPRv2 (52961), pMD2.G (12259) were from Addgene. Cu standard (SC194-500), Fe standard (SI124-500), TMT mass tagging kit 6-plex (90061), RIPA lysis buffer (89900), Epredia™ SuperFrost™ Plus Slides (6776214) were from Thermo Fisher Scientific. Ammonium acetate (A637), formaldehyde (NC9603389), trace metal-grade nitric acid (A509) were from Fisher Scientific. Anti-CTR1 (Proteintech, 67221-1-Ig), anti-CAD antibody (Proteintech, 16617-1-AP), anti-ATP7A (Santa Cruz, sc-376467), anti-giantin (abcam, ab80864), HRP-conjugated secondary antibody (ab205719), qPCR Master Mix (Ambion, 4472908), Mn standard (Acros Organics, 196111000), EIPA (Cayman, 14406), $H_2S$ gas (Linde/Cryogenic Gases, custom order), pLentiCRISPRv2-OR7G3 (gift from the Mootha lab, MGH), elesclomol (gift from the Gohil lab, TAMU), high pH reversed-phase peptide fractionation kit (Pierce, 84868), silicon nitride ($SiN_x$) membrane windows (Norcada, NX5150D and NX5150E), Zn standard (Inorganic Ventures, CGZN1), PhenoPlate (Revvity, 6057302), Kapton film (Cole-Parmer, UX0457595), Cytoseal mounting medium (Epredia, 8310-16), and Histoclear (National Diagnostics, HS-200) were from the specified vendors.

### Animal studies

The generation of $Sqor^{fl}/^{fl}$ mice and the intestinal epithelial–specific SQOR KO $Villin^{Cre}$ $SQOR^{fl/fl}$ mice as well as the antibiotic treatment have been described[3] and all animal protocols were approved by the Institutional Animal Care and Use Committee at the University of Michigan. Mice (all ~10 weeks old; without antibiotics: 1 male and 2 females per genotype; with antibiotics: 1 male and 2 females per genotype) were euthanized with $CO_2$ followed by secondary cervical dislocation. Colon was dissected, rinsed and the distal portion was used for ICP-MS analysis. The proximal colon was Swiss-rolled, embedded in optimal cutting temperature (OCT) compound, and used for X-ray fluorescence microscopy (XFM), laser ablation ICP-MS (LA-ICP-MS) and haematoxylin and eosin (H&E) staining.

### Cell culture

All cells were maintained at 37°C in a humidified atmosphere containing 5% $CO_2$ and passaged every 3 to 5 days. HT-29 cells were cultured in RPMI 1640 supplemented with 10% FBS and 1% penicillin/streptomycin. The shRNA KD lines (*SLC31A1*, *CTR1*, *ATP7A* and *CAD*) and CRISPR KO (*OR7G3*, *SLC30A1*/*ZNT1*) HT-29 lines were maintained with the addition of 1 μg/ml puromycin. EA.hy926 and HEK293 143B, H9c2 parental and Ctr1 KO cells were cultured in DMEM with 10% FBS and 1% penicillin/streptomycin.

For acute treatment, freshly prepared $Na_2S$ stock solution was added at 0 h (100 μM) and 1 h (100 μM), for a total exposure of 2 h. Cells were maintained in a separate incubator to prevent cross-contamination of untreated cells with volatile sulfide. For chronic sulfide exposure, cells were cultured in a custom-built sulfide growth chamber[4] in which $H_2S$ gas (25 or 100 ppm) was mixed with 5% $CO_2$ in air for normoxic conditions or with 93% $N_2$, 5% $CO_2$, and 2% $O_2$ for hypoxic conditions. For recovery experiments, cells were exposed to chronic sulfide for 24 h, after which the medium was changed, and cells were transferred to a 5% $CO_2$ incubator with humidified ambient air for the indicated times. When used, ES-Cu (in DMSO) and $ZnSO_4$ (in Milli-Q water for qPCR analysis) were supplemented 2 h before, and EIPA (in DMSO), BCS (in Milli-Q water) and $ZnSO_4$ (for ICP-MS analysis) were added at the start of sulfide treatment.

### Cell proliferation assay

HT-29 cells ($6\times10^5$ cells) and HEK293 ($2\times10^5$ cells) were seeded in 6-well plates containing either 2 mL complete RPMI + 25 mM HEPES or DMEM medium. The next day (≈16 h), the medium was changed (2 mL) and cells were grown ± 100 ppm $H_2S$ for 24 h. Then, the cells were harvested or moved to a regular $CO_2$ incubator (21% $O_2$). The medium was changed every 24 h (2 mL). For cell counting, cells were washed once in 1x PBS, trypsinized and resuspended in 1xPBS + 10 mM EDTA. Cell suspensions were mixed 1:1 with trypan blue and counted using a Cellometer Auto T4 Brightfield Cell Counter. Dead cells were excluded from the live cell count analysis.

### Generation of stable KD lines

Stable KDs of *SLC31A1/CTR1, ATP7A* and *CAD* were generated using pLKO.1 lentiviral vectors with shRNA sequences listed in Supplementary Table 4; control lines were generated using a non-targeting scrambled shRNA vector. CRISPR KD of *SLC30A1/ZNT1* and an *OR7G3*-

targeting control plasmid were based on pLentiCRISPRv2. The pLentiCRISPRv2-OR7G3 plasmid was from the Mootha laboratory and the *SLC30A1*/*ZNT1* KD lines were generated using guide RNA (sgRNA) sequences cloned into the pLentiCRISPRv2 vector. Briefly, the following oligonucleotides were used for ZNT1:sgRNA3 5'-CACCGGATCCGAGCCGAGGTAATGG-3' and sgRNA4 5'-CACCGGCCGCAGCCATGGGGTGTTG-3'. Each oligonucleotide pair was annealed and ligated into BsmBI-digested pLentiCRISPRv2. Insertion was verified by Sanger sequencing using the hU6-F (5'-GAGGGCCTATTTCCCATGATT-3') and LKO.1_5 (5'-GACTATCATATGCTTACCGT-3') primers.

Lentivirus batches were prepared by seeding 2 × $10^6$ HEK293 cells per 6-cm dish in 4 mL of complete DMEM. At ~70% confluency, cells were transfected with 750 ng psPAX2, 250 ng pMD2.G, and 1 μg of the lentiviral transfer vector using 12 μL lipofectamine 3000 and 4 μL P3000 reagent in serum-free Opti-MEM, following the manufacturer's protocol. After 48 h, the supernatant containing the lentiviral particles was collected, filtered through a 0.45 μm membrane, and used immediately for transduction. HT-29 cells (~70% confluency in 6-cm plates) were transduced by adding 1 mL of filtered viral supernatant to 3 mL complete RPMI 1640. After 24 h, fresh medium containing 1 μg/mL puromycin was used to replace the transduction medium Following 4 to 5 days of selection, KD efficiency was confirmed by Western blot and/or ICP-MS analysis.

### Western blot analysis

HT-29 cells (2.4 × $10^6$ per 6-cm dish or 3.6 × $10^6$ cells per 10-cm dish) were cultured for 16-20 h in 4 or 10 mL of complete RPMI 1640 with 25 mM HEPES, respectively. Then, the medium was changed (8 or 20 mL) and cells were grown for an additional 24 h ± 100 ppm $H_2S$. For ATP7A and CAD analysis, cells were washed in ice-cold 1× PBS, scraped into pre-weighed tubes, resuspended in 4 μL/mg wet weight of lysis buffer (20 mM HEPES, pH 7.5, 25 mM KCl, 0.5% NP-40, 10 μL/mL protease inhibitor cocktail), and stored at -80 °C. For CTR1 analysis, pellets were resuspended in 4 volumes ($^w/_v$) of detergent-free lysis buffer (20 mM HEPES, pH 7.5, 25 mM KCl, 10 µL/mL protease inhibitor cocktail) and stored at -80 °C. ATP7A and CAD KD lysates were prepared by three freeze-thaw cycles (1 min at 37 °C; 10 min on dry ice) and clarified at 13,000 × *g* for 5 min at 4 °C. CTR1 membrane fractions were obtained by centrifugation at 21,000 × *g* for 15 min at 4 °C, weighed, and solubilized in 3 µL/mg wet weight of lysis buffer with 1% Triton X-100 for 30 min on ice, with 10 s vortexing every 10 min, then clarified at 21,000 × *g* for 5 min at 4 °C.

Total protein was quantified by the Bradford assay using BSA as a standard. For CTR1, BSA standards were prepared in the 1% Triton X-100 lysis buffer. ATP7A and CAD KD samples were normalized to 2.5 µg/µL in 4× Laemmli buffer (200 mM Tris-HCl, pH 6.8, 40% glycerol, 4% SDS, 200 mM DTT) and heated at 65 °C for 10 min; CTR1 samples were prepared at 3.5 µg/µL and heated at 37 °C for 15 min. Equal amounts of protein were resolved on precast 12% or 4-20% SDS-PAGE gels and transferred to ethanol-activated 0.2 µm PVDF membranes for 30 min using the transfer buffer supplied with the Trans-Blot Turbo Transfer Kit. Membranes were blocked in 5% non-fat dry milk in TBS-T (1× TBS, 0.1% Tween 20) for 1 h at room temperature and incubated overnight at 4 °C with primary antibodies against ATP7A (1:100), CAD (1:600) or CTR1 (1:1,000). After 3X TBS-T washes, membranes were incubated with HRP-conjugated secondary antibody (1:10,000) for 1-2 h at room temperature and developed with ECL substrate. Total protein was assessed by Ponceau S staining of the membrane. Densitometry was performed in Image Lab (Bio-Rad) and target signals were normalized to total protein.

### RNA-Seq and qRT-PCR analysis

HT-29 cells were seeded at 1.2 × $10^6$ cells per well in six-well plates in 2 mL of complete RPMI 1640 and cultured for 16 h. The medium was changed (2 mL) and $Na_2S$ was added at 0 h (100 µM) and 1 h (100 µM). After 2 h, cells were washed twice in 1× PBS, lysed in TRIzol, and stored at -80 °C until extraction. RNA was extracted using TRIzol, according to the manufacturer's instruction, washed in 75% ethanol, and resuspended in nuclease-free water. RNA integrity was confirmed by agarose gel electrophoresis. Library construction and sequencing were performed by Novogene Corporation (Sacramento, CA) on an Illumina NovaSeq platform.

FASTQC (v0.11.9) was used to verify sequencing data quality, and then the fastq files were provided as input to a primary analysis pipeline, originally published by Rogers et al.,[5] consisting of the following steps. STAR aligner (v2.7.5a),[6] with option --twoPassMode Basic specified, was used to align the reads to human genome reference hg38 with ENCODE blacklisted regions masked (accession ENCFF356LFX) and GENCODE annotation version 33. Reads that mapped equally well to both the mitochondrial genome and the nuclear genome were assigned to chrM using a custom script (prioritize_chrM_over_numts.py (see Rogers et al. 2023)[5]. Duplicate reads were collapsed using Picard MarkDuplicates as provided in the GATK package (v4.1.5.0-9-g227bef6-SNAPSHOT).[7] Then, reads were assigned to genes using the featureCounts program provided in the Subread package (v2.0.1)[8] with the options -t exon, -p, -M, and --primary. Finally, the featureCounts results were compiled into a count matrix and combined with metadata using the AnnData package (v0.7.5) in Python (v3.7.12). DESeq2 (v1.30.1)[9] running in R (v4.0.5) was used to test for differentially expressed genes between the $H_2S$ and control samples after filtering out genes with fewer than 10 reads.

For chronic sulfide exposure, $3.6 \times 10^6$ HT-29 cells were seeded in 10-cm dish in 10 mL of RPMI 1640 with 25 mM HEPES and cultured for 20 h. The medium was changed (20 mL) and cells were exposed to chronic sulfide (100 ppm $H_2S$, 24 h). Total RNA was extracted using the TRIzol reagent. RNA yield and purity were determined using a NanoDrop spectrophotometer, and samples were stored at -80°C. Complementary DNA (cDNA) was synthesized from 1 µg of total RNA using M-MLV reverse transcriptase and random hexamers. Quantitative PCR was performed using SYBR™ Select Master Mix and a QuantStudio Real-Time PCR System. Primer sequences are listed in Supplementary Table 5 and all samples were run in duplicate or triplicate. Relative gene expression was calculated using the $2^{-\Delta\Delta Ct}$ method. To ensure robust normalization, the geometric mean of three independent reference genes (*GUSB*, *TBP*, and *POLR2A*) was utilized as the internal control, as described previously.[10] The RNA-Seq data have been deposited in the GEO repository with the dataset identifier GSE341519.

#### TMT proteomics analysis

HT-29 cells were seeded in triplicates at 7.2 × $10^6$ cells per 10 cm dish in 10 mL complete RPMI 1640 medium with 25 mM HEPES. The next day (~16 h), the medium was replaced with 20 mL of fresh medium and the cells were exposed to ± 100 ppm $H_2S$ for 24 h. Cells were washed once in 1x PBS, trypsinized and resuspended in RIPA lysis buffer (4 µL/mg wet weight) with 10 µL/mL protease inhibitor cocktail. Protein was quantified by the Bradford assay and samples (75 µg of protein per condition) were stored at -80°C and submitted to the Proteomics Resource Facility at the University of Michigan.

Briefly, upon reduction (5 mM DTT, for 30 min at 45°C) and alkylation (15 mM 2-chloroacetamide, for 30 min at room temperature) of cysteines in samples, the proteins were precipitated by adding 6 volumes of ice-cold acetone followed by overnight incubation at -20°C, centrifuged and then, the pellet was allowed to air dry. The pellet was resuspended in 0.1 M triethylammonium bicarbonate and digested overnight (~16 h) with trypsin/Lys-C mix (1:25 protease:protein; Promega) at 37°C with constant mixing in a thermomixer. The TMT 6-plex reagents (TMT mass tagging kit 6-plex) were dissolved in 41 µL anhydrous acetonitrile and labeling was performed by transferring the entire digest to the TMT reagent vial and incubating at room temperature for 1 h. The reaction was quenched by adding 8 µl 5% hydroxylamine and incubated for 15 min. The labeled samples were mixed, and dried using a vacufuge. An offline fractionation of the combined sample (~200 µg) into 8 fractions was performed using high pH reversed-phase peptide fractionation kit according to the manufacturer's protocol. Fractions were dried and reconstituted in 9 µL 0.1% formic acid/2% acetonitrile in preparation for LC-MS/MS analysis. Samples (3 controls and 3 sulfide treated) were labeled with TMT channel 126-128 and 129-131.

For superior quantitation accuracy, multinotch-MS3[11] was employed, which minimizes the reporter ion ratio distortion resulting from fragmentation of co-isolated peptides during MS analysis. An Orbitrap Ascend Tribrid spectrometer equipped with FAIMS source (Thermo Fisher Scientific) and Vanquish Neo UHPLC was used for data acquisition. Two μL of each sample was resolved on an Easy-Spray PepMap Neo column (75 μm i.d. x 50 cm; Thermo Scientific) at a flow-rate of 300 nL/min using 0.1% formic acid/acetonitrile gradient system (3-19% acetonitrile in 72 min;19--29% acetonitrile in 28 min; 29-41% in 20 min followed by 10 min column wash at 95% acetonitrile and re-equilibration) and directly sprayed onto the MS using EasySpray source (Thermo Fisher Scientific). FAIMS source was operated in standard resolution mode, with a nitrogen gas flow of 4.2 L/min, and inner and outer electrode temperature of 100°C and dispersion voltage or -5000 V. Two compensation voltages (CVs) of -45 and -65 V, 1.5 seconds per CV, were employed to select ions that enter the MS for MS1 scan and MS/MS cycles. The MS was set to collect MS1 scan (Orbitrap; 400-1600 m/z; 120K resolution; AGC target of 100%; max IT in Auto) following which precursor ions with charge states of 2-6 were isolated by quadrupole mass filter at 0.7 m/z width and fragmented by collision induced dissociation in ion trap (NCE 30%; normalized AGC target of 100%; max IT 35 ms). For multinotch-MS3, top 10 precursors from each MS2 were fragmented by HCD followed by Orbitrap analysis (NCE 55; 45K resolution; normalized AGC target of 200%; max IT 200 ms, 100-500 m/z scan range).

Data were analyzed using Proteome Discoverer (v3.0; Thermo Fisher). MS2 spectra were searched against SwissProt human protein database (20352 entries; *Homo sapiens* (sp_canonical TaxID=9606) (v2023-09-13); downloaded on 11/02/2023) using the following search parameters: MS1 and MS2 tolerance were set to 10 ppm and 0.6 Da, respectively; carbamidomethylation of cysteines (57.02146 Da) and TMT labeling of lysine and N-termini of peptides (229.16293 Da) were considered static modifications; oxidation of methionine (15.9949 Da) and deamidation of asparagine and glutamine (0.98401 Da) were considered variable. The proteins and peptides identified were filtered to retain only those that passed ≤1% FDR threshold. Quantitation was performed using high-quality MS3 spectra (average signal-to-noise ratio of 10 and <50% isolation interference). The mass spectrometry proteomics data have been deposited in the ProteomeXchange Consortium via the PRIDE partner repository[12] with the dataset identifier PXD079024.

**GC-MS analysis**

HT-29 (empty vector, and with either cytoplasmic or mitochondrially targeted *Lb*NOX) were seeded at a density of 3.6 x $10^6$ cells/10-cm dish in 10 mL complete RPMI+HEPES and cultured ± 100 ppm $H_2S$ for 24 h. Then, the medium was aspirated and cells were washed twice with 5 mL chilled (4°C) 1x PBS. Metabolites were quenched by the addition of 1 mL ice-cold 80% methanol (pre-chilled on dry ice for 30 min). Plates were incubated on dry ice for 10 min, after which the cells were scraped and the extract was collected. The suspension was centrifuged at 16,000 x *g* for 10 min at 4°C. The supernatant was transferred to a fresh tube, and the total volume recovered was recorded for normalization to cell counts. Samples were stored at -80°C until analysis.

GC-MS analysis was performed on an Agilent 7890B GC system equipped with a DB-5MS capillary column (30 m, 0.25 mm i.d., 0.25 µm-phase thickness; Agilent J & W Scientific), connected to an Agilent 5977B Mass Spectrometer operating under ionization by electron impact (EI) at 70 eV. Helium flow was maintained at 1 mL/ min. The source temperature was maintained at 230 °C, the MS quad temperature at 150 °C, the interface temperature at 280 °C, and the inlet temperature at 250 °C.

The samples were evaporated to dryness using a vacuum concentrator. Dried samples were methoxylamine hydrochloride tert-butyldimethylsilyl (MOX-TBDMS) derivatized and analyzed by GC-MS, as described previously.[13] Carbamoyl aspartate eluted with a retention time of 26.85 min and was identified using the characteristic m/z 443 fragment ion. Absolute amounts were determined using a calibration curve generated with authentic carbamoyl aspartate samples of known concentration. All mass spectra were recorded in single ion monitoring (SIM) mode with 4 ms dwell time on each ion and integrated to determine total ion counts, as described previously.[14] Data were normalized to cell count and expressed as intracellular molar concentrations calculated by assuming a cell volume of $2.4 \times 10^{-12}$ L.

### Immunofluorescence and confocal microscopy

HT-29 scrambled and ATP7A KD cells were seeded at 2,000 cells/well in 40 µL of complete RPMI with 25 mM HEPES in 384-well imaging plates (PhenoPlates). Glass surfaces were coated with poly-L-lysine for 1 h at 37 °C and 5% $CO_2$, then washed once with 1× PBS prior to seeding. Following chronic sulfide exposure (± 100 ppm $H_2S$, 24 h), the culture medium was aspirated, and cells were washed once with room-temperature 1x PBS. Cells were fixed in 4% paraformaldehyde in 1x PBS for 10 min at room temperature in the dark, washed three times in 1x PBS, permeabilized with 0.25% Triton X-100 in 1× PBS for 10 min, and blocked in 5% BSA with 0.3% Triton X-100 in 1× PBS for 30 min. Primary antibodies (anti-ATP7A 1:50; anti-giantin 1:100), diluted in 1x PBS-T (1% BSA, 0.3% Triton X-100), were incubated overnight at 4 °C. Following three washes with 1x PBS-T, cells were incubated with Alexa Fluor-conjugated secondary antibodies (anti-mouse 488 and anti-rabbit 555; both 1:1 000) for 1 h at room temperature in the dark. Nuclei and cell membranes were counterstained with Hoechst (1:1 000) and CellMask (1:10 000) in 1x PBS for at least 1 h. Samples were stored in 1x PBS with sodium azide at 4°C until further use. Imaging was performed on a Yokogawa CellVoyager 8000 dual spinning-disk confocal high-content microscope with a 40 x water immersion objective and a 50 µm pinhole. For each field of view, 10 z-slices were acquired across 10 µm (1 µm step) and collapsed into a maximum-intensity projection. For image analysis, the Python package spaCR (v.1.0.0)[15] was used to segment cells and nuclei, and capture intensity and morphological measurements from each channel. Co-localization between ATP7A and the Golgi marker Giantin was quantified using Pearson's correlation coefficient (PCC).

#### Transmission electron microscopy (TEM)

Endosomes were analyzed in previously collected TEM images of HT-29 cells cultured ± 100 ppm $H_2S$ for 24 h.[16] The images were blinded prior to manual annotation. 976 endosomes from 39 control cells and 854 endosomes from 43 $H_2S$-exposed cells were counted.

#### Total reflection X-ray fluorescence (TXRF) analysis

HT-29 ($7.2 \times 10^6$) cells were seeded in triplicates per 10 cm dish in 10 mL complete RPMI 1640 medium. The next day (~16 h), the medium was changed (20 mL) and cells were exposed to $H_2S$ (either acute or chronic) as described above. Cells were harvested by trypsinization, resuspended in complete medium and centrifuged at 1,600 x *g* for 3 min at 4°C. The resulting pellet was washed once with 1 mL ice-cold 1x PBS with 10 mM ethylenediamine tetraacetic acid (EDTA) to remove extracellular metals, followed by a final wash and resuspension in 1x PBS. For cell counting, aliquots of the suspensions were mixed 1:1 with trypan blue and counted using a Cellometer Auto T4 Brightfield Cell Counter (Nexcelom). For TXRF analysis, aliquots of $4\times10^6$ cells (live + dead) were centrifuged, and the pellets were snap-frozen in liquid nitrogen and lyophilized overnight at -84°C under vacuum. Samples were stored at -80°C.

TXRF analysis was performed as previously described.[17] Briefly, frozen cells pellets were thawed on ice and resuspended in 5 μL of 1x PBS spiked with 1 ppm gallium (Ga). A 100 ppm Ga stock in 5% nitric acid ($HNO_3$), which was prepared from a master stock of 10,000 ppm Ga in 5% $HNO_3$ (Ricca Chemical Company), was added to give a final concentration of 1 ppm in 1 x PBS. Silicone solution (10 μL) (SERVA) was spotted onto the center of a quartz sample disk and dried by heating on a hot plate set at 120 °C for 5 min. Following this, 2 μLs of the cell suspension to be analyzed was spotted onto the center of the siliconized quartz disc and dried by heating at 120 °C for 5 min to form a thin film. Multi-elemental analysis (Fe, Cu, and Zn) was performed using a Bruker S2 PICOFOX TXRF spectrophotometer operating at 50 kV and 600 μA. Spectra were collected over a 1500s sampling time. Data were processed using the S2 Picofox spectral deconvolution software and metal concentrations were normalized to cell number and expressed as a fold change relative to untreated control cells.

### ICP-MS analysis

Cells were seeded at low (~10%) confluency in 10-cm plates with 10 mL of culture medium as follows: $3.6\times10^6$ (HT-29 in RPMI+ HEPES), $2.5\times10^6$ (HEK293, 143B, complete DMEM) or $1\times10^6$ (EA.hy926, complete DMEM). The next day, the medium was changed (20 mL) and cells were incubated ± 100 ppm $H_2S$ (24 h). Cells were harvested as described above (under TXRF analysis). Cell pellets were digested in 15 mL tubes in trace metal-grade nitric acid (50 μL/$4\text{x}10^6$ cells, stock 67-70%) and boiled at 80°C for 2 h. Then, $H_2O_2$ (25 μL/$4\times10^6$, stock 30% ($^w/_w$)) was added, and samples were incubated at 85 °C for an additional hour in loosely capped tubes. Samples were allowed to cool to room temperature and centrifuged for 1 min at 3000 x *g*. A blank sample with $HNO_3$ and $H_2O_2$ was processed in parallel. All samples were subsequently diluted to a final nitric acid concentration of 4% ($^v/_v$) with Milli-Q $H_2O$. Metal standards (Fe, Cu and Zn) were prepared in 4% nitric acid. Samples and standards were analyzed on a NexION 2000 ICP-MS (PerkinElmer) with yttrium as an internal standard. Cu and Zn were monitored in the standard collision mode while $^{57}Fe$ was monitored in Helium-Kinetic Energy Discrimination mode. Metal amount was normalized to the total (live and dead) cell count.

For ICP-MS analysis of tissue samples, frozen distal colon sections were weighed and digested in 15 mL tubes with trace metal-grade nitric acid (10 μl/mg wet tissue) at 80 °C for 2 h. An empty tube devoid of tissue sample was processed in parallel as a blank. Samples were diluted to 4% $HNO_3$ ($^v/_v$) with Milli-Q water, analyzed by ICP-MS and Cu amount was normalized to tissue wet weight.

### XFM analysis

*XFM analysis of cells*. HT-29 and HEK293 cells were imaged on ultra-thin silicon nitride ($SiN_x$) membrane windows, which were placed in 6-well plates and coated with 20 μL of sterile 0.01% poly-L-lysine for 30 min at 37°C to promote cell adhesion. After removing excess poly-L-lysine, the windows were seeded with a 20 μL droplet of cell suspension ($1 \times 10^5$ cells/mL HT-29 or $2.5 \times 10^4$ cells/mL HEK293). Cells were allowed to adhere for 10 min at 37°C before the wells were gently flooded with an additional 2 mL of the respective cell suspension. Cells were then cultured ± 100 ppm $H_2S$ for 24 h, at 2% or 21% $O_2$ and with or without recovery in normal air. Then, the culture medium was removed and replaced with 4% paraformaldehyde in 1x PBS for 30 min at 37°C to fix cells. To remove extracellular salts and media components, the $SiN_x$ windows were subjected to a series of 5-min washes: 3X in 1x PBS, 3X in freshly prepared 0.1 M isotonic ammonium acetate, and 3X in Milli-Q water. Windows were transferred to a filter paper and allowed to air-dry completely at room temperature. Samples were stored at room temperature until further analysis. XFM was performed at beamline HXN-3ID at NSLSII, Brookhaven National Laboratory (Figures 1 and 3 and Supplementary Figure 5) and at the Bionanoprobe on beamline 2IDD at APS, Argonne National Laboratory (Supplementary Figures 2 and 6). $SiN_x$ windows were mounted and scanned at an incident X-ray energy of 10 keV. The monochromatic X-ray beam was focused using Fresnel Zone plates. A silicon drift detector positioned 90 degrees to the incident beam was used to collect XRF spectra. Maps were acquired with a spatial resolution of 100-200 nm (Figure 1), 64-125 nm (Figure 3 and Supplementary Figure 5) or 100-125 nm (Supplementary Figures 2 and 5) and dwell times of 200-250 ms (Figure 1)/100-150 ms (Figure 3 and Supplementary Figure 5) and 150 ms (Supplementary Figures 2 and 5). Elemental maps were obtained by fitting the XRF spectra at each pixel and quantified using pyXRF software[18] or XRF-MAPS software (https://github.com/AdvancedPhotonSource/XRF-Maps). Absolute elemental areal densities (grams/cm$^2$) were determined by calibrating the XRF intensity with thin-film standards, measured under the same conditions as the samples.

*XFM analysis of tissue.* Tissue blocks stored at -80°C were moved to -20°C 24 h prior to sectioning. Blocks were mounted in a cryostat (Leica Biosystems CM3050s) with a chamber temperature of -20°C and sample temperature of -22°C. Sections were mounted onto Epredia™ SuperFrost™ Plus (positively charged) slides for laser ablation imaging and H&E staining or 8 μm Kapton film for XFM. XFM sections (20 μm) and corresponding H&E sections (10 μm) were separated by 40 μm, while the laser ablation imaging sections (10 μm) and corresponding H&E sections (10 μm) were adjacent.

XFM was performed at 5-ID Submicron Resolution X-ray Beamline at the National Synchrotron Light Source-II (Brookhaven National Laboratory, Upton, NY). Samples were shipped on dry ice and stored at -80°C until they were imaged. A Si(111) double-crystal monochromator was used to select for 10 keV x-rays, which were focused to a 500 x 500 nm spot size (FWHM) using a pair of Kirkpatrick-Baez mirrors (JTEC Corporation). Scans were performed at 500 nm step sizes with 25 ms dwell time. XFM data were collected using a 7-element Vortex ME-7 Silicon Drift Detector (Hitachi, USA), normalized to the incident x-ray beam intensity measured using an upstream ion chamber filled with nitrogen, then fitted and visualized using the M-BLANK software package.[19] Briefly, XFM data were blank corrected using the measured spectrum of an empty Kapton substrate and the residual XFM emissions were modeled and fitted using modified Gaussians.[20, 21] Emission energies, fluorescence yields, and branching ratios[22, 23] were referenced using the *CXRO X-ray Data Booklet*[24] and the xraylib[25] program, with branching ratios empirically optimized during parameter creation. Once calculated, peak shapes were held constant and used in linear least squares fitting of the spectral data in M-BLANK. Fitted counts for each element were converted to areal density (μg/cm) by comparison to a thin film reference standard (AXO standard RF11-200-S3235) containing Ca (2.67 μg/cm), Fe (0.44 μg/cm), and Cu (0.23 μg/cm). Calibration values for elements not present were either interpolated or extrapolated. A full list of instrument setup and parameters has been described previously.[26] Areas of interest for XFM analysis were determined by examining adjacent sections with LA-ICP-MS and H&E staining.

### LA ICP-MS analysis

Colon sections were scanned using a Bioimage 266 (Elemental Scientific Lasers) LA system with a 10 μm spot size, 60% laser power, and 125 Hz scan rate. The ablation system was coupled to a Tofwerk custom ICP-TOF-MS for elemental analysis. To quantify elemental content of the colon samples, 10 weight percent gelatin standards doped with either a liquid standard mixture (Inorganic Ventures custom standard IV-74434: 1000 μg/mL As, B, Ca, Cd, Co, Cr, Cu, Fe, K, Mg, Mn, Mo, Na, Ni, Pb, Se, V, Zn) or sodium phosphate mono- or dibasic (Sigma S0751 and S9763, respectively) were prepared with a thickness matching the samples. An aliquot of each gelatin standard was measured using an ICP-MS (Agilent 8900 ICP triple quadrupole) to obtain the concentration of each element in each gelatin standard. Standard concentrations: 0, 12.5, 25, and 50 ppm IV-74434 and 2000, 4000, and 8000 ppm NaP. Areas of interest for quantification were selected as described previously.[27, 28] Briefly, an iterative thresholding approach was applied to the phosphorus map. Initially, pixels exceeding an intensity of 10% the maximum were labeled as sample and the remaining set was refined by repeatedly excluding values above the mean plus two standard deviations until the population became self-consistent.

The mean Cu content in the ROI of each tissue was normalized by the mean P content to control for cell content. For H&E staining of adjacent sections a previously described protocol was used.[29] Briefly, slides were fixed in 4% paraformaldehyde for 5 min, then rinsed in Millipore water for 1 min. The samples were stained with hematoxylin for 3 min, placed in defining media for 20 sec, and Blue Buffer for 40 sec, with each step separated by a 1-min water wash. After the last water wash, the slide was dehydrated with 95% ethanol for 40 sec then stained with eosin for 12 sec. The slides were then washed with two 40 sec 95% ethanol washes, followed by three 2-min 100% ethanol washes. Finally, the samples were cleared with two 3-min Histoclear rinses, mounted with Cytoseal mounting medium, and a coverslip was placed over the sample. Samples were imaged on a Zeiss Axioscan 7 slide scanner at 10x magnification.

#### XAS and EXAFS analysis

HT-29 ($5\times10^6$ cells per 10-cm plate) were seeded in quadruplicates in 10 mL complete RPMI 1640 with HEPES. The next day, the medium was changed (20 mL) and cells were incubated ± 100 ppm $H_2S$ for 24 h and harvested as described under “TXRF analysis”. After the final 1xPBS wash, cell pellets were weighed and resuspended in 1xPBS with 50% glycerol to a final glycerol concentration of 17%. Cell suspensions were loaded into cuvettes sealed with 25 μm-thick Kapton tape, flash frozen, and stored in liquid nitrogen until analysis.

XAS spectra were measured at the Stanford Synchrotron Radiation Light source on beamlines 9-3. A Si(220) double crystal monochromator, oriented at Φ=0, was used for energy selection at the Cu K-edge. The harmonic rejection was provided by a rhodium-coated mirror. The beam was focused to ~250 μm using an XOS polycapillary optics. Sample surface was scanned to locate high Cu-concentration spots for data collection. The elastic and inelastic scattering signal from the sample was reduced using a Ni filter and Soller slits. Samples were maintained at 10 K in a helium atmosphere during data collection using an Oxford liquid helium cryostat. Spectra were collected in the fluorescence mode using a multi-element Canberra Ge detector, and a Cu foil reference spectrum was collected in the transmission mode simultaneously with each scan for energy calibration. The spectrum of CuO and $Cu_2O$ were acquired from XASDB,[30, 31] and the collection of other reference spectra has been described before.[32, 33]

During data processing, channels with large background signals from ice diffraction were removed. The remaining channels were then averaged in ATHENA.[34] Data were normalized and calibrated by setting the edge energy (first maximum in first derivative) of the Cu foil to 8980.3 eV. The normalized spectra were imported into PySpline[35] for post-edge background subtraction. EXAFS was extracted by setting $E_0$ to 9000 eV and fitting the background using a 4-region spline (with polynomial orders 2, 3, 3, and 3). Fitting of $k^3$-weighted EXAFS was performed using EXAFSPAK,[36] and the theoretical phase and amplitude of scattering paths were generated using FEFF7.[37] The starting geometry of the models was adopted from DFT geometry optimization.

### Data preparation and statistical analysis

Unless noted otherwise in the figure legends, each data point is from an independent experiment whereas n = 1 for animal tissue studies, represents data from a single mouse. Data were analyzed and graphs were generated with R v4.5.3 (base, ggplot2, tidyverse), RStudio 2026.01.1+403. Data are presented as dot plots with mean (line) or line graphs with means and error bars showing standard deviation (SD) or violin plots with median and mean. For all data sets where p-values are indicated, normal distribution (Shapiro-Wilk, stats::shapiro.wilk) and homogeneity of variances (Levene, car::leveneTest) were tested. For comparisons between two groups, an unpaired two-sided Two-sample t-test (stats::t.test) was performed for equal variance, a two-sided Welch Two sample t-test for unequal variance (stats::t.test) or a two-sided Wilcoxon rank sum test (stats::wilcox.test) for data sets with outliers.

**Supplementary Table 3. Cu K-edge EXAFS curve-fitting analysis.**

| Sample | Coordination/path | $R$ (Å)[a] | $\sigma^2$ (Å$^2$×10$^5$) | $\Delta E_0$ (eV) | $F^b$ |
|---|---|---|---|---|---|
| HT-29 cells grown with chronic $H_2S$ | 3 Cu-O/N | 1.96 | 543 | -7.66 | 0.320 |
| | 1 Cu-O/N | 2.52 | 854 | | |
| | 5 Cu-C | 2.95 | 718 | | |
| | 6 Cu-C-O | 3.21 | 134 | | |
| | 4 Cu-C | 3.72 | 1395 | | |
| | 1 Cu-Cu | 4.05 | 1333 | | |
| Fit without Cu-Cu path | 3 Cu-O/N | 1.97 | 542 | -7.52 | 0.338 |
| | 1 Cu-O/N | 2.52 | 857 | | |
| | 5 Cu-C | 2.95 | 703 | | |
| | 6 Cu-C-O | 3.21 | 139 | | |
| | 4 Cu-C | 3.73 | 1751 | | |
| Fit using four paths | 3 Cu-O/N | 1.96 | 542 | -8.13 | 0.349 |
| | 1 Cu-O/N | 2.51 | 823 | | |
| | 5 Cu-C | 2.95 | 745 | | |
| | 6 Cu-C-O | 3.19 | 171 | | |

[a]The estimated standard deviations for the bond lengths are on the order of ±0.02 Å. [b]The error is given by $[\Sigma k^6(\chi_{exptl}-\chi_{calcd})^2/\Sigma k^6\chi_{exptl}^2]^{1/2}$. The $S_0^2$ factor was set at 1. The number of independent parameters in the EXAFS spectrum is calculated as $N_{ind}=2\Delta k\Delta R/\pi+2=23$. The best fit includes 13 independent parameters. The disorder in the Cu-Cu path is high, and the EXAFS fit alone cannot be used to make a definite assignment of a Cu-Cu component. Additional fits show the first shell parameters are robust and less affected by the outer shells.

**Supplementary Table 4**. pLKO.1 shRNA vectors.

| Gene | Product code | Target Sequence |
|---|---|---|
| SLC31A1/CTR1 | TRCN0000043349 | 5'-GATGCCTATGACCTTCTACTT-3' |
| SLC31A1/CTR1 | TRCN0000043350 | 5'-GCGTAAGTCACAAGTCAGCAT-3' |
| ATP7A | TRCN0000043173 | 5'-CCATTCATGTACTAGCACTAT-3' |
| ATP7A | TRCN0000043174 | 5'-GCTGTATTAGTAGCAGTTGAT-3' |
| ATP7A | TRCN0000043177 | 5'-CCTCTTGGTATGGATTGTAAT-3' |
| CAD | TRCN0000045908 | 5'-GCTCCGAAAGATGGGATATAA-3' |
| CAD | TRCN0000045910 | 5'-CGAATCCAGAAGGAACGATTT-3' |
| CAD | TRCN0000045912 | 5'-CCCAGATGAAATGGATGAGTT-3' |

**Supplementary Table 5.** Oligonucleotides used for qPCR

| Oligonucleotide | Sequence |
|---|---|
| MT1A_F | 5'-AGAGTGCAAATGCACCTCCTGC-3' |
| MT1A_R | 5'-CGGACATCAGGCACAGCAGCT-3' |
| MT1G_F | 5'-AGAGTGCAAATGCACCTCCTGC-3' |
| MT1G_R | 5'-TTGTACTTGGGAGCAGGGCTGT-3' |
| MT1X_F | 5'-AGAGTGCAAATGCACCTCCTGC-3' |
| MT1X_R | 5'-TGTCCTGGCATCAGGCACAGC-3' |
| MT2A_F | 5'-GAGTGCAAATGCACTTCGTGCAA-3' |
| MT2A_R | 5'-GCGTTCTTTACATCTGGGAGCG-3' |
| GUSB_F | 5'-CTGTCACCAAGAGCCAGTTCCT-3' |
| GUSB_R | 5'-GGTTGAAGTCCTTCACCAGCAG-3' |
| TBP_F | 5'-TGTATCCACAGTGAATCTTGGTTG-3' |
| TBP_R | 5'-GGTTCGTGGCTCTCTTATCCTC-3' |
| POLR2A_F | 5'-GAGAGCGTTGAGTTCCAGAACC-3' |
| POLR2A_R | 5'-TGGATGTGTGCGTTGCTCAGCA-3' |

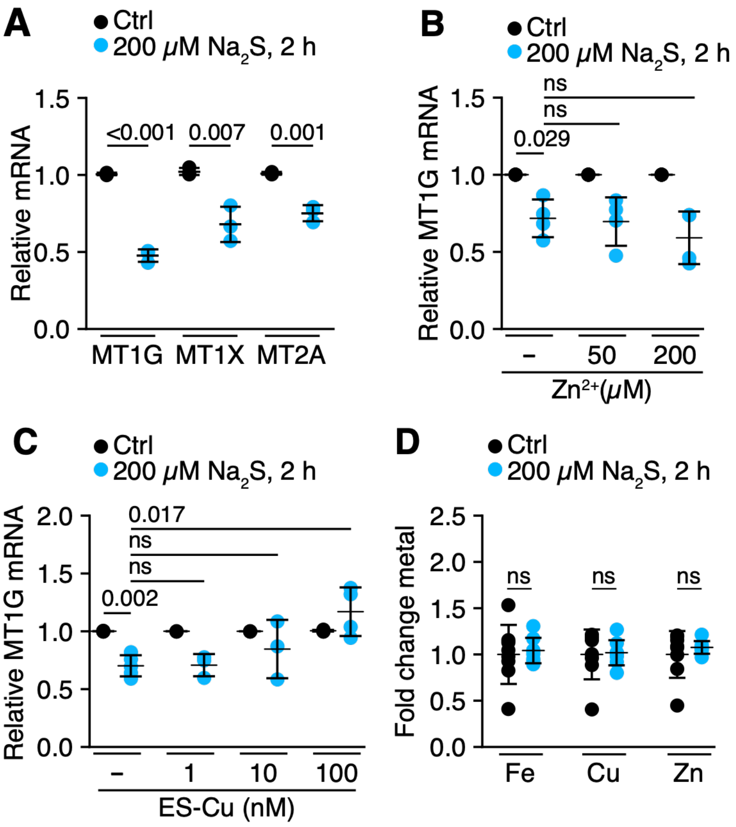


**Supplementary Figure 1. Acute $H_2S$ exposure decreases MT but not intracellular Cu. A.** qPCR confirmed MT1G, MT1X and MT2A, are significantly less abundant in response to acute $H_2S$ (200 µM, 2h), n = 3**. B.** Zn (50 or 200 µM) did not prevent MT1G mRNA downregulation in response to acute $H_2S$ (n = 4). Sulfide-treated samples are normalized to the respective controls (0, 50 and 200 µM $Zn^{2+}$). Basal MT1G mRNA increased 29- and 571-fold in response to 50 and 200 µM $Zn^{2+}$, respectively, as expected **C**. High (100 nM) but not low (1 or 10 nM) Cu-elesclomol prevented MT1G downregulation in response to acute $H_2S$ treatment (n=3-5). Sulfide-treated samples are normalized to the respective controls (0, 1, 10 and 100 nM ES-Cu). Basal MT1G mRNA remained unchanged in response to 1 and 10 nM ES-Cu but increased 38-fold in response 100 nM ES-Cu. **D**. Significant differences in total Fe, Cu or Zn levels were not observed by TXRF spectroscopy in response to acute $H_2S$ treatment (n = 2, with each data point representing a technical replicate). Data show mean ± SD.

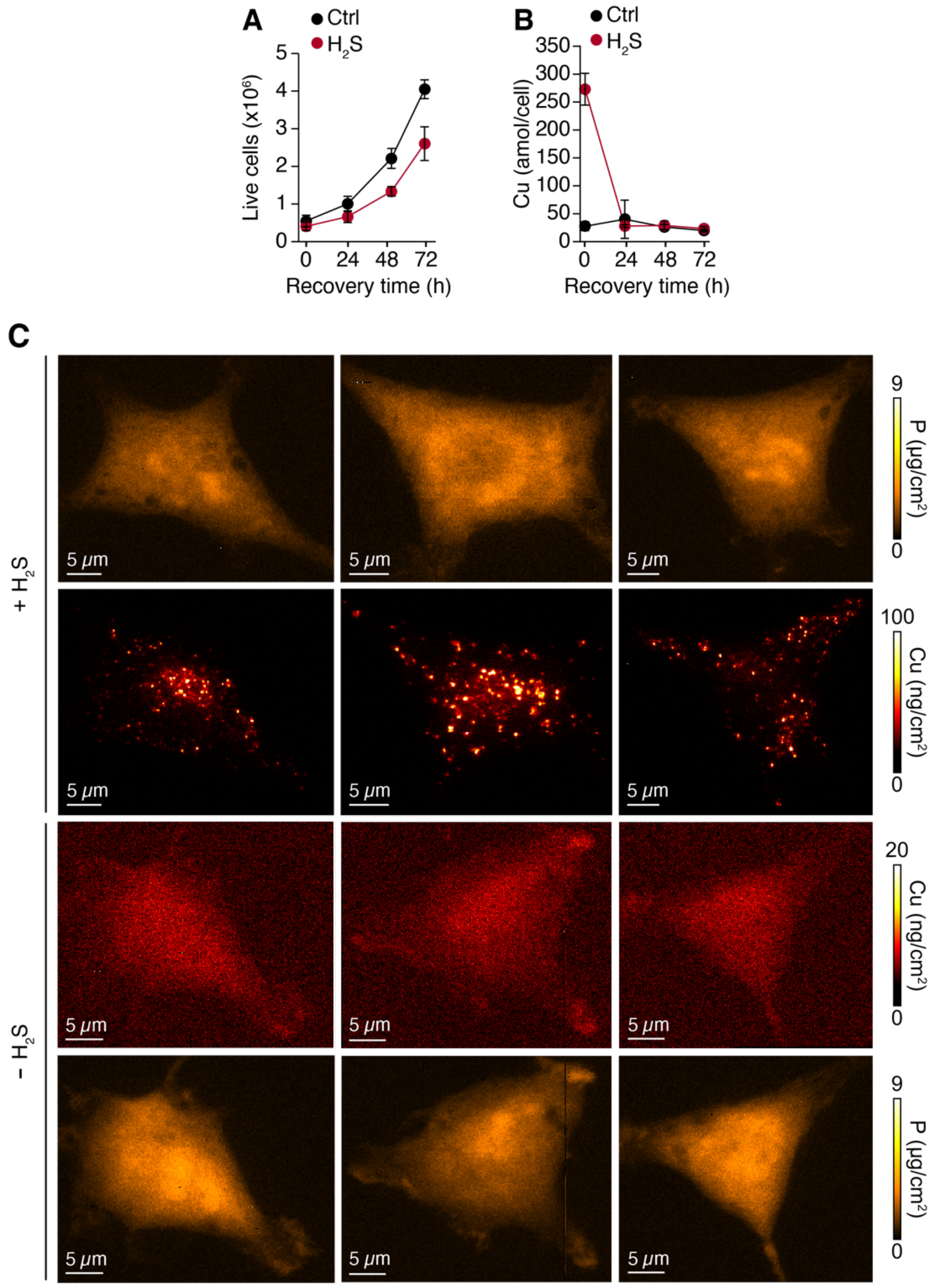


**Supplementary Figure 2**. **Sulfide induces Cu puncta formation in HEK293 cells**. **A,B**. HEK293 cells resume proliferation (A, n=3) and normalize Cu to basal levels within 24 h (B, n=5) following sulfide withdrawal. **C**. Representative XFM images of three HEK293 cells exposed to chronic $H_2S$ at 21% $O_2$, showing Cu puncta (*upper*) compared to three untreated cells (*lower*). A more sensitive scale has been used in the lower panel to visualize Cu in untreated cells, which have a lower concentration. The corresponding phosphorus (P) images are shown above (+$H_2S$) and below (-$H_2S$) the respective Cu images and serves to outline the cell.

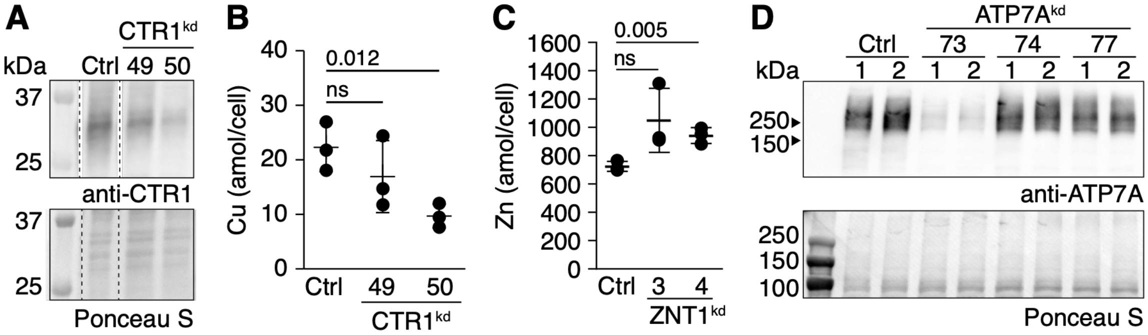


**Supplementary Figure 3. Validation of CTR1, ATP7A and ZNT1 KDs in HT-29 cells. A, B.** Western blot analysis (A) and basal Cu levels (B, n=3) in control (Ctrl) versus CTR1 KDs using shRNA #49 and #50 **C**. Effect of ZNT1 KD using CRISPR sgRNA3 and sgRNA4 on basal Zn levels as determined by ICP-MS (n=1-3). **D**. ATP7A KD with shRNA 73 but not 74 or 77 led to a significant decrease in protein levels. In A and D, Ponceau S staining was used to assess equal loading.

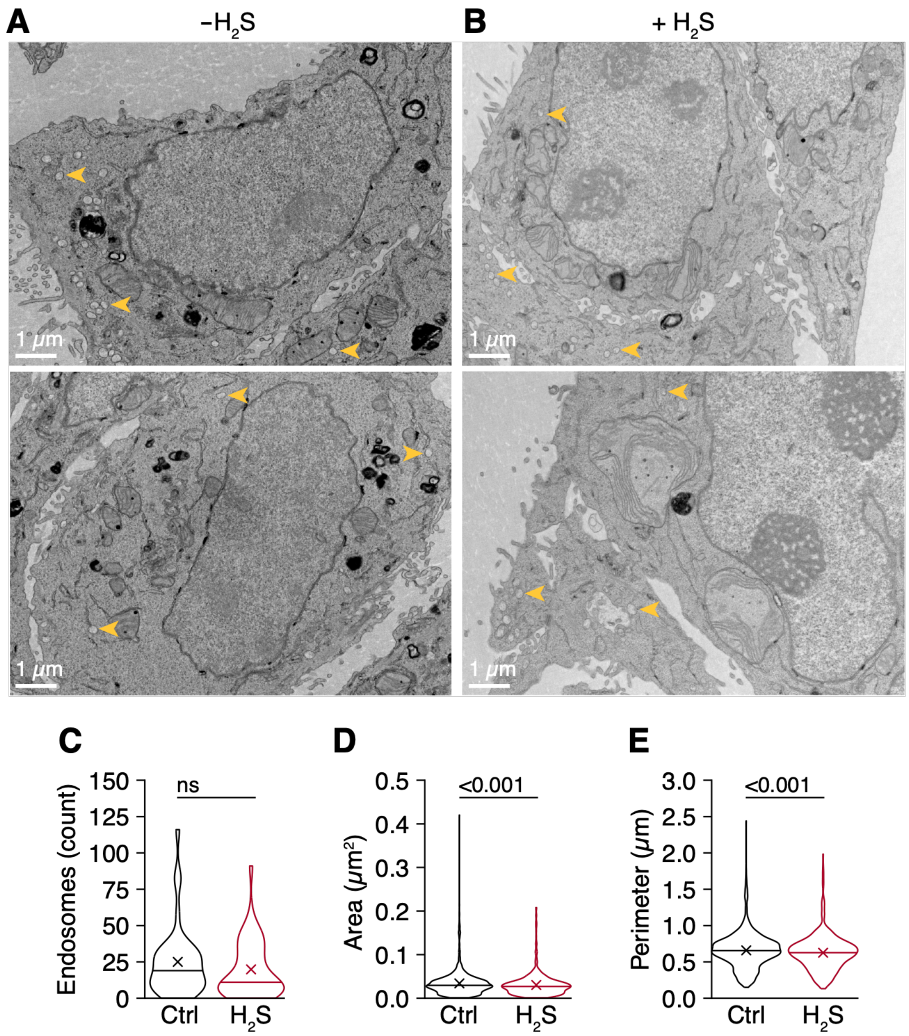


**Supplementary Figure 4. Transmission electron microscopy analysis of endosomes in HT-29 cells. A**, **B**. Two representative electron micrographs comparing cells grown with or without chronic $H_2S$ exposure. Yellow arrow heads are pointing to representative endosomes. **C-E**. Quantification reveals that the abundance of endosomes in untreated versus $H_2S$ exposed cells is comparable (C) and although the difference in area (D) and perimeter (E) were statistically significant they were marginally different. Violin plot shows median (line) and mean (cross), 39 (ctrl) and 43 ($H_2S$) cells/condition.

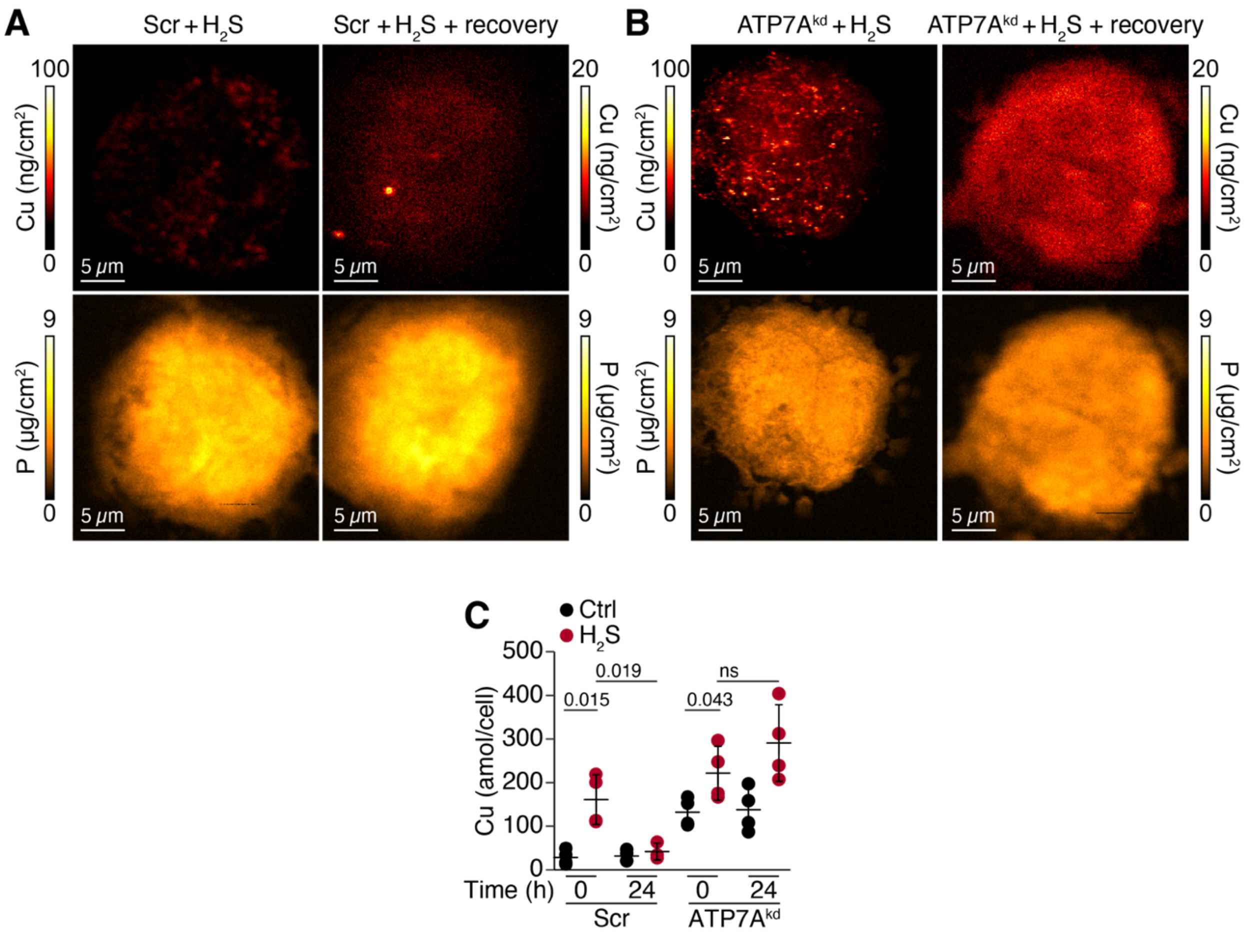


**Supplementary Figure 5. ATP7A is required for Cu export, but not dissolution of Cu puncta after sulfide withdrawal. A, B.** Following sulfide withdrawal after chronic $H_2S$ exposure (100 ppm, 24 h), scrambled control (Scr) and ATP7A KD cells showed loss of Cu puncta. Representative Cu (*top*) and the corresponding phosphorus (P) images are shown (*below*). Note the difference in scales for Cu in sulfide-treated and sulfide-treated + recovered cells. **C**. ICP-MS analysis of whole cell Cu in HT-29 cells following sulfide withdrawal (n=3 or 4 independent experiments). The 0 h time point refers to the start of the experiment (i.e. after 24 h of chronic sulfide exposure) and is identical to the data shown in Fig. 3F; 24 h refers to the time after $H_2S$ withdrawal.

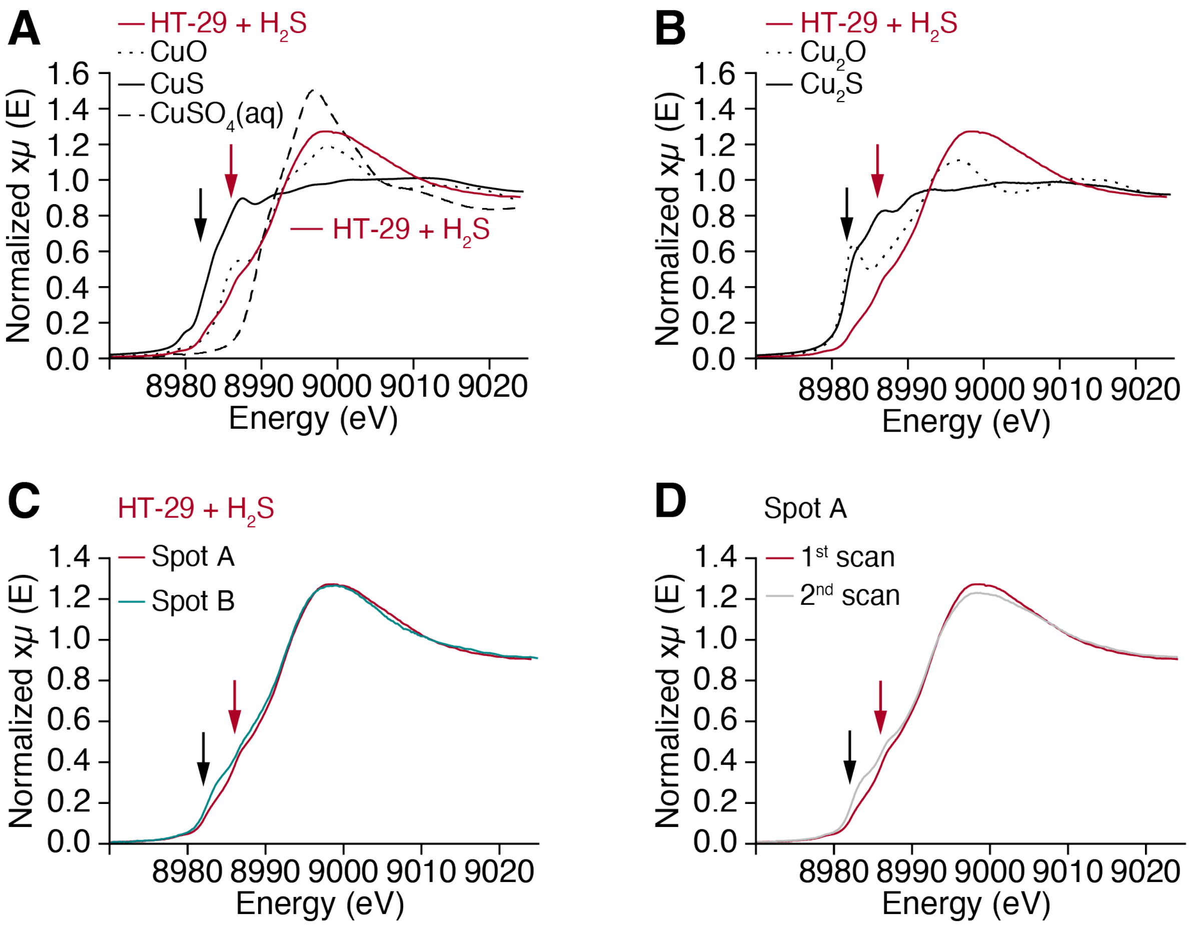


**Supplementary Figure 6. Predominance of $Cu^{2+}$ and the effects of spot-to-spot variation and sample photoreduction on Cu K-edge XAS. A, B.** Comparison of Cu K-edge XAS collected from HT-29 cells and comparison to $Cu^{2+}$ (A) and $Cu^{1+}$ (B) compounds. **C,** Cu K-edge XAS collected from two spots with high Cu concentrations, showing variation in the relative $Cu^{1+}$ and $Cu^{2+}$ fractions. **D.** First and second XAS scans collected at spot A, showing beam-induced photoreduction of $Cu^{2+}$. Photoreduction increases the rising edge intensity at ~8983 eV (black arrow) while decreasing the white line intensity and EXAFS oscillation amplitude. The EXAFS from the second scan at spot A was used for analysis because it provided the highest signal-to-noise ratio. However, the reduced EXAFS amplitude may lead to slight overestimation of the Debye-Waller factors for the first shell ligands. Black and red arrows indicated the rising edge intensities at ~8983 eV and ~8986 eV for $Cu^{1+}$ and $Cu^{2+}$, respectively.

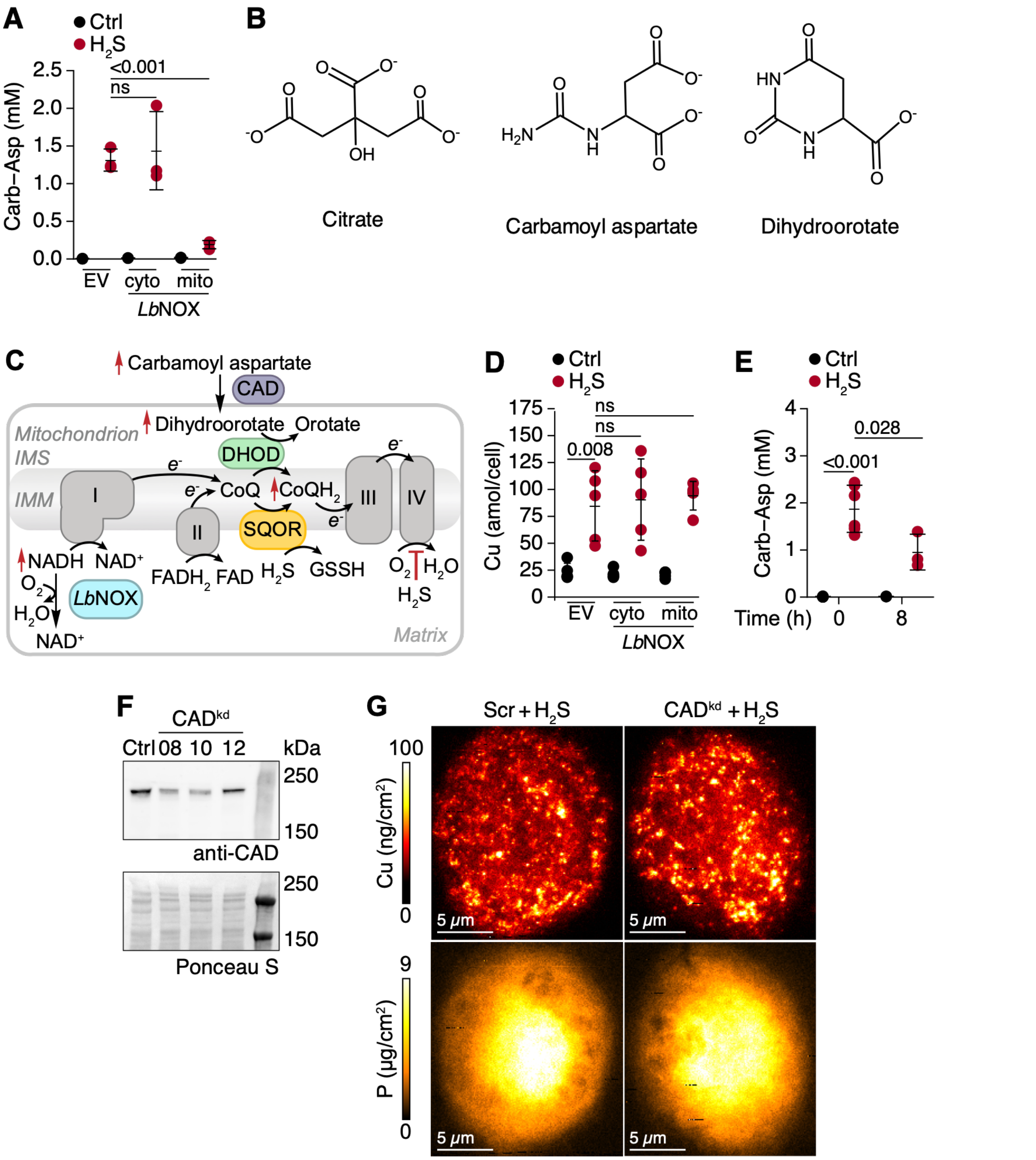


**Supplementary Figure 7. $H_2S$-induced Cu accumulation is independent of carbamoyl aspartate. A.** Expression of mitochondrial, but not cytosolic *Lb*NOX decreases carbamoyl-aspartate under chronic $H_2S$ exposure (100 ppm, 24 h) (n=3). **B.** Structures of the indicated metabolites. **C.** Scheme showing $H_2S$-dependent inhibition of complex IV induces a reductive shift that leads to increased levels of carbamoyl aspartate and dihydroorotate. **D.** Neither expression of mitochondrial nor cytosolic *Lb*NOX decreases intracellular Cu under chronic $H_2S$ exposure (n=5). **E.** Carbamoyl aspartate levels decrease significantly after 8 h of sulfide withdrawal following chronic $H_2S$ (n=3-6). **F**. Western blot analysis in scrambled control (Ctrl) versus CAD KDs using shRNA #8, 10 and 12. Ponceau S stained membrane is shown below for equal loading. **G**. Representative XFM images of scrambled versus CAD KD (shRNA #10) HT-29 cells exposed to chronic $H_2S$ at 21% $O_2$, showing Cu puncta (*upper*) and the corresponding phosphorus (P) images (*lower*) which serve to outline the cell.
